# A hyperparameter-randomized ensemble approach for robust clustering across diverse datasets

**DOI:** 10.1101/2023.12.18.571953

**Authors:** Sarah M. Goggin, Eli R. Zunder

## Abstract

Clustering analysis is widely used to group objects by similarity, but for complex datasets such as those produced by single-cell analysis, the currently available clustering methods are limited by accuracy, robustness, ease of use, and interpretability. To address these limitations, we developed an ensemble clustering method with hyperparameter randomization that outperforms other methods across a broad range of single-cell and synthetic datasets, without the need for manual hyperparameter selection. In addition to hard cluster labels, it also outputs soft cluster memberships to characterize continuum-like regions and per cell overlap scores to quantify the uncertainty in cluster assignment. We demonstrate the improved clustering interpretability from these features by tracing the intermediate stages between handwritten digits in the MNIST dataset, and between tanycyte subpopulations in the hypothalamus. This approach improves the quality of clustering and subsequent downstream analyses for single-cell datasets, and may also prove useful in other fields of data analysis.

## Introduction

Clustering is widely used for exploratory data analysis across diverse fields, where it is applied to identify dataset grouping structures in an unsupervised manner. In particular, clustering has become a workhorse tool for single-cell analysis, enabling the identification and characterization of cell populations that share similar molecular profiles within heterogeneous biological samples^1^. The output of clustering analysis is often used for direct comparison of biological samples, to identify changes in the abundance or molecular state of specific cell populations. Furthermore, clustering output is frequently carried forward into additional downstream analyses such as cell type classification or trajectory analysis^2,3,4^. Therefore, the accuracy and reproducibility of clustering partitions is important for the quality of single-cell analysis. This importance has motivated the development of hundreds^5^ of clustering methods with a variety of algorithmic strategies, but there are still important shortcomings in all of these methods which reduce their effectiveness.

An ideal clustering method for single-cell analysis would satisfy the following requirements:

1. Operate without the need for human input such as hyperparameter tuning. The vast majority of existing methods require selection and optimization of hyperparameters, which can significantly impact clustering quality ^6–9^. Manual hyperparameter tuning is time-consuming and relies subjectively on human intuition about which groupings appear correct ^10^. Automated methods have been proposed to overcome this limitation, but many are computationally inefficient, and all are biased by the criteria used for optimization ^9,11,12,13^.
2. Perform well across diverse single-cell datasets from different tissues and across multiple measurement modalities such as single-cell/single-nucleus RNA sequencing (scRNA-seq and snRNA-seq), single-cell Assay for Transposase-Accessible Chromatin sequencing (scATAC-seq), flow cytometry, mass cytometry, and multiplexed imaging analysis such as high-content fluorescence imaging, imaging mass cytometry (IMC), multiplexed ion beam imaging (MIBI), and multiplexed error-robust fluorescence in situ hybridization (MERSCOPE). Generalizability is a concern in existing methods; many clustering methods perform well on gold-standard single-cell datasets, but do not generalize well to datasets from other tissue types or from other single-cell analysis modalities which may have different or more complex distributions or structural properties ^7,8,9,10,14^.
3. Produce stable and consistent partitions that are robust to random sampling and minor perturbations. Existing methods do not reliably produce robust partitions when applied to complex, high dimensional single-cell datasets. Meaningfully different results can be produced with different hyperparameter combinations ^8^, slight perturbations of a dataset ^10,14^, or even when an identical dataset and hyperparameters are run multiple times due to randomization steps in most clustering algorithms (Supplementary Fig. 1a,b).
4. Capture and describe the wide variety of discrete and continuous grouping structures present in single-cell datasets ^15,16^. Most existing methods implement hard clustering, which assumes a data structure with discrete, well-separated groups, but is unable to characterize overlap or continuity between groups. Alternative computational methods for trajectory inference can better capture specific types of continuum-like processes such as cell differentiation in single-cell datasets, but these methods make a different set of assumptions about data structure that can be equally restrictive.
5. Quantify uncertainty at the levels of individual data points and clusters. There are many scenarios where clustering can provide useful information, but a single optimal solution to the clustering task either does not exist or cannot be determined ^17^. In many cases, there is additionally no known ground truth that could define what a correct solution might look like. Therefore, measures of uncertainty are crucial to assess the reliability and aid interpretability of clustering results before using them as inputs for downstream analytical methods or for purposes such as hypothesis development or orthogonal validation of results.
6. Scale to analyze large single-cell datasets with millions of cells. While many of the most commonly used methods are scalable, several that have been developed to address these key challenges for clustering have done so at the expense of scalability. Methods that improve on these other challenges can only be realistically impactful if they can produce results for the large dataset sizes that are becoming increasingly commonplace.

Recently developed clustering methods have made progress towards some of these goals. Ensemble and consensus methods represent a promising approach to improve clustering robustness by combining information from multiple diverse partitions ^18–25^. Fuzzy and soft clustering allow data points to belong to multiple clusters, and can therefore be used to provide a more complete description of both continuous and discrete data structures ^26,27^. There are several methods that provide measures of stability or uncertainty at the cluster level ^9,11,24,28^, but cell-level measures of uncertainty are rarely provided in single-cell methods ^29,30^. However, none of these approaches have been able to incorporate all of the six key features described above.

To address this need for a single method that performs robustly across diverse datasets with no hyperparameter tuning and transparently communicates uncertainty, we developed a clustering algorithm that applies <u>E</u>n<u>S</u>emble <u>C</u>lustering with <u>H</u>yperparameter <u>R</u>andomization (ESCHR). This algorithm requires no human input due to hyperparameter randomization, which explores a wide range of data subspaces that contribute to the final consensus clustering step. Our implementation of ESCHR in Python (https://github.com/zunderlab/eschr) can be used as a self-contained framework for clustering, or it can be integrated into commonly used single-cell analysis pipelines such as the scverse ecosystem ^31^. To evaluate this new method, we performed extensive benchmarking tests, which demonstrated that ESCHR outperforms both general clustering methods and the most widely used clustering methods for single-cell analysis ^24,32–34^, both in terms of accuracy on synthetic datasets with a known “ground truth,” and in terms of robustness on real single-cell datasets encompassing diverse tissues (bone marrow, pancreas, developing and adult brain), organisms (mouse, human), cell numbers (from hundreds to millions), and measurement techniques (single-cell RNA sequencing, mass cytometry, flow cytometry).

After benchmarking for accuracy and robustness, we applied ESCHR clustering to two complex real-world datasets -first to the MNIST dataset ^35^, a commonly used example for machine learning image analysis, and then in the single cell context to investigate the relationships between tanycyte populations in the hypothalamus, which have been previously shown to display spatial and molecular-level continuity between subtypes ^36–40^. In both of these exploratory analyses, the soft cluster assignments and overlap scoring from ESCHR were used to identify regions of low confidence cluster assignments corresponding to transitional overlap between clusters and map the key feature transitions that define these regions.

## Results

### Overview of ESCHR clustering

To develop a robust and scalable clustering method for analysis of single-cell datasets, we employed an ensemble and consensus approach, which has been shown to improve robustness across many domains of machine learning ^21,41,42,43,44,45^. This approach consists of two main steps: 1) generate a set of base partitions, referred to as the ensemble, and 2) use this ensemble to generate a final consensus partition. The graph-based Leiden community detection method ^46^ was selected as a base algorithm to generate the clustering ensemble, because it is widely used for single-cell analysis, and is efficiently implemented to be scalable for large datasets ^3^.

A key element of successful consensus approaches is generating sufficient diversity in the ensemble ^21,42,43^. To generate this diversity, ESCHR randomizes four hyperparameters for each base partition: subsampling percentage, number of nearest neighbors, distance metric, and Leiden resolution. Within a given base partition, ESCHR first selects a subsampling percentage by random sampling from a gaussian distribution with μ scaled to dataset size (within 30-90%), and then extracts the specified subset of data from the full dataset. Next, ESCHR randomly selects values for the number of nearest neighbors (15-150) and the distance metric (euclidean or cosine) and uses these to build a k-nearest neighbors (kNN) graph for the extracted subset of data. Finally, ESCHR performs Leiden community detection on this kNN graph using a randomly selected value for the required resolution-determining hyperparameter (0.25-1.75). The ranges for randomization of these hyperparameters were optimized empirically (Supplementary Fig. 2a-h and Methods). This subsampling and randomization scheme is used to produce diversity amongst each of the different base partitions (Fig. 1a). This diversity provides many different views of the dataset, and the full ensemble of these views provides a more comprehensive picture of the dataset grouping structure, which is less likely to be influenced by the stochastic variations present in any single view, including the full unsampled dataset. In addition to generating ensemble diversity, this hyperparameter randomization approach is what enables ESCHR to operate without the need for hyperparameter tuning at this first stage of the algorithm.

**Figure 1:**
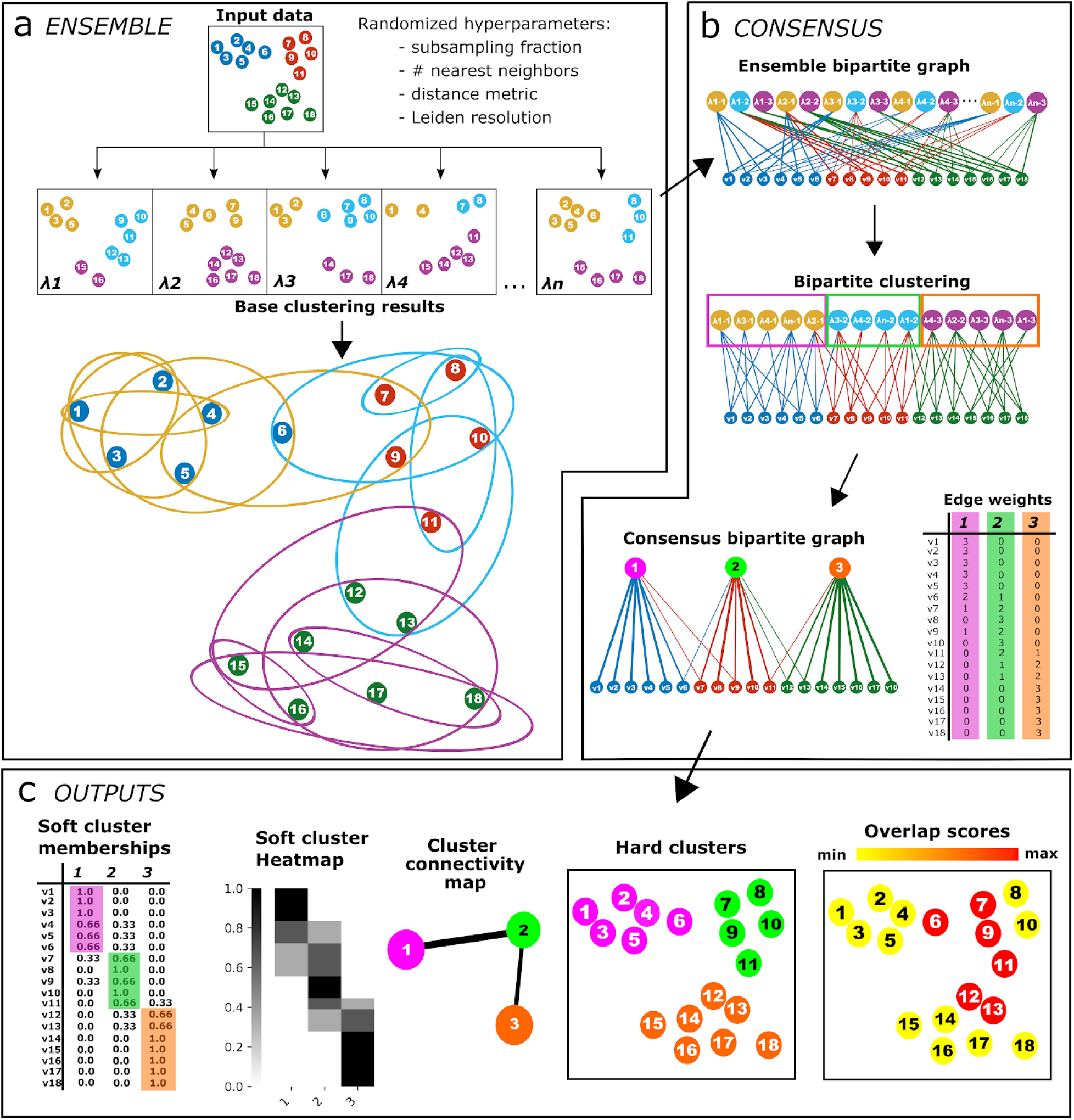
ESCHR framework overview. (A) Starting from a preprocessed input dataset, ESCHR performs ensemble clustering using randomized hyperparameters to obtain a set of base partitions. This set of base partitions is represented using a bipartite graph where one type of node consists of all data points and one type of node consists of all clusters from all base partitions and edges exist between data points and each base cluster they were assigned to throughout the ensemble. (B) Leiden bipartite clustering is performed on the ensemble bipartite graph. Base clusters are collapsed into their assigned consensus clusters obtained through the bipartite clustering and edge weights are summed such that each data point now has a weighted edge to each consensus cluster representing the number of base clusters it had been assigned to the were then collapsed into that consensus cluster. (C) Soft cluster memberships are obtained by scaling edge weights between 0 and 1, and can then be visualized directly in heatmap form and used to generate hard cluster assignments, per-data point overlap scores, and cluster connectivity maps.

After generating a diverse ensemble of base partitions, ESCHR applies a bipartite graph clustering approach to obtain the final consensus partition. First, the base partitions are assembled into a bipartite graph, where cells are represented by one set of vertices, base clusters are represented as a second set of vertices, and each cell is connected by an edge to each of the base clusters it was assigned to throughout the ensemble (Fig. 1b). Next, ESCHR applies bipartite community detection to obtain the final consensus partition (Fig. 1b) ^47^.

Bipartite community detection is applied here instead of more common consensus approaches that suffer from information loss ^48^. To remain hyperparameter-free without the need for human intervention in this consensus stage of the algorithm, ESCHR performs internal hyperparameter selection to determine the optimal resolution for the final consensus clustering step by selecting the medoid from a range of resolutions (Supplementary Fig. 3a-d). After obtaining the final consensus partition, ESCHR converts the ensemble bipartite graph to a final weighted bipartite graph by collapsing all base partition cluster nodes assigned to the same consensus cluster into a single node. Cells are then connected to these consensus cluster nodes by edges with weights representing the number of times each cell was assigned to any of the base partition clusters that were collapsed into a given consensus cluster (Fig. 1b). These raw membership values are then normalized to obtain proportional soft cluster memberships, and hard cluster labels are assigned as the consensus cluster in which a cell has the highest proportional membership (Fig. 1c).

While many analysis strategies for single-cell datasets require hard clustering labels, these by definition cannot convey whether a cell is at the borderline between multiple clusters or located firmly in the center of a single cluster. Hard clusters also do not provide any insight into potential continuity between clusters. Using the soft cluster memberships derived from the weighted consensus bipartite graph, ESCHR provides several additional outputs beyond hard cluster assignments that enable more comprehensive characterization of the grouping structures within a dataset. Firstly, soft cluster memberships can be directly visualized in heatmap form to identify areas of cluster overlap at the single-cell level (Fig. 1c). Importantly, these soft membership heatmap visualizations can serve as complements or even alternatives to the widely used but also widely misinterpreted ^49^ stochastic embedding methods (i.e. UMAP ^50^, t-SNE ^51^) for visualizing the complex relational structures within single-cell datasets. ESCHR also produces an Overlap Score for every object, derived from its soft cluster membership, which quantifies regions of higher and lower certainty in hard cluster assignment (Fig. 1c). Finally, ESCHR produces a cluster-level map of the continuity structure within a dataset by using the soft cluster memberships to calculate a corrected-for-chance measure of the connectivity between each pair of hard clusters (Fig. 1c and Methods).

### ESCHR soft clustering and overlap scores capture diverse structural characteristics and quantify uncertainty in cluster assignments

We first sought to examine how ESCHR overlap scores and soft clustering could enable effective and informative analysis for datasets containing complex combinations of continuity and discreteness, and how these results compared to a wide range of alternative clustering methods used for single-cell analysis or general purpose clustering (Supplementary Table 1 and Methods). For this analysis, we desired to use datasets with known true labels, so we assembled a collection of 20 structurally diverse synthetic datasets that consist of randomly generated gaussian distributions varying in number of objects (5000 or 10000), number of features (20,40,50,60), number of clusters (3,8,15,20), cluster sizes, cluster standard deviations, cluster overlap, and cluster anisotropy (Supplementary Fig. 4a-d). In our analysis, we refer to these individual gaussian distributions as ground truth clusters.

We selected two illustrative examples from these synthetic datasets, each of which represent different challenges for traditional hard clustering approaches. The first dataset, with 3 gaussian distributions and 5000 objects total, is prone to overclustering with the default hyperparameters of most clustering algorithms, but ESCHR and SC3 were able to identify the correct number of clusters (Figs. 2a, S5a). The overclustering primarily occurred in ground truth cluster 2, which has a high degree of overlap with ground truth cluster 1. This level of uncertainty, related to the difficulty in clustering, is reflected in the ESCHR overlap scores (Fig. 2b). We validated this observation using the F1 score, which combines the concerns of both precision and recall into a single score and can be calculated individually for each true cluster ^52^. All clustering methods were able to accurately capture the true cluster 0, which has the lowest overlap with other clusters, but true clusters 1 and 2 were more challenging (Fig. 2c). In addition to quantifying this level of uncertainty and clustering difficulty via the per cell Overlap Scores, ESCHR also provides information about which clusters overlap, and to what extent, as visualized in the soft cluster membership heatmap, which reveals substantial overlap between ESCHR clusters 0 and 2 (corresponding to true labels 1 and 2) (Fig. 2d).

**Figure 2:**
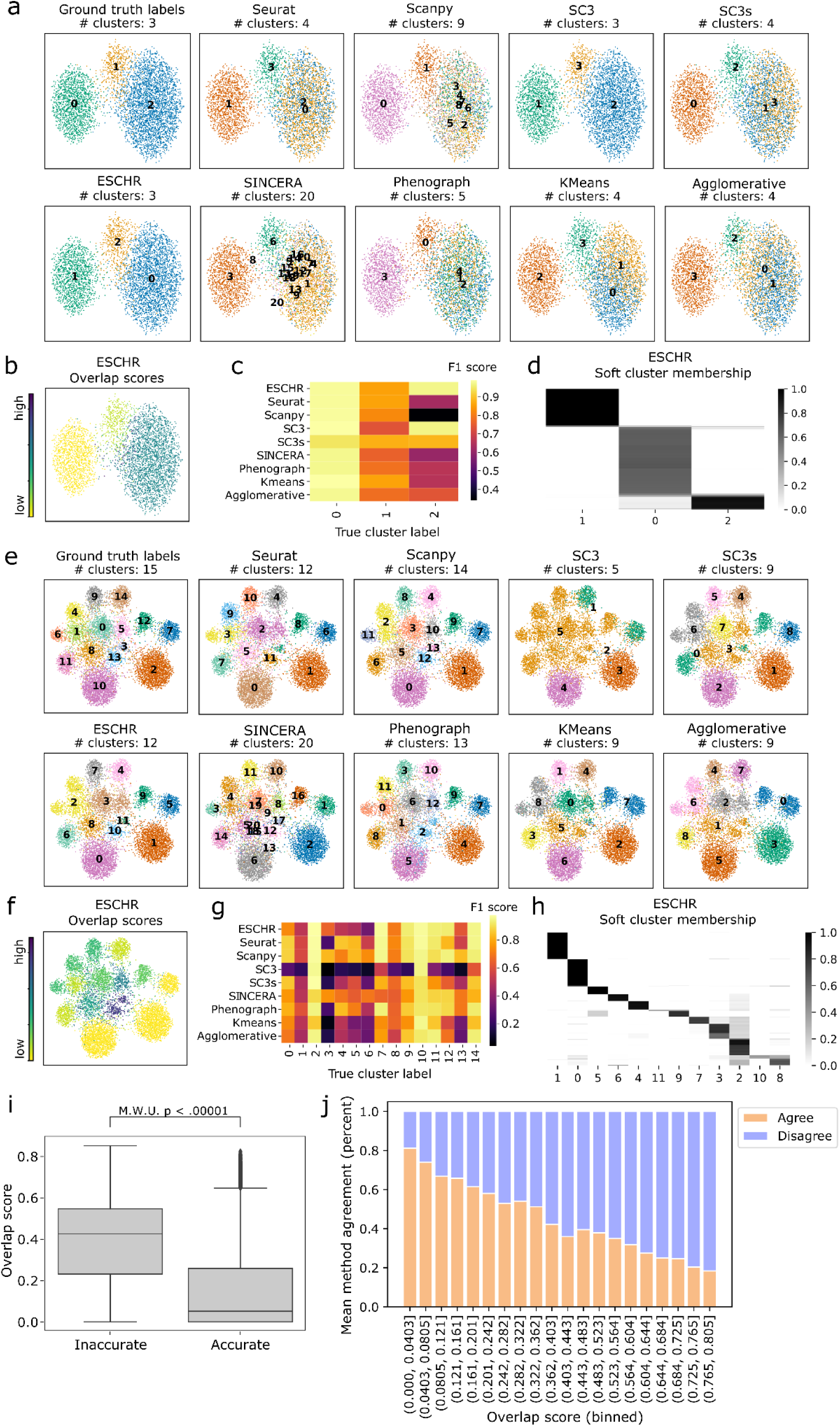
Visualization of ESCHR clustering and overlap scores compared to other clustering methods. (A) UMAP visualizations of ground truth cluster labels and hard cluster assignments from ESCHR and selected comparison methods. Points are colored by cluster ID. (B) UMAP visualization with points colored by ESCHR overlap score. (C) F1 score calculated per method for each ground truth cluster separately. (D) Heatmap visualization of ESCHR soft cluster memberships. (E) UMAP visualizations of ground truth cluster labels and hard cluster assignments from ESCHR and selected comparison methods. Points are colored by cluster ID. (F) UMAP visualization with points colored by ESCHR overlap score. (G) F1 score calculated per method for each ground truth cluster separately. (H) Heatmap visualization of ESCHR soft cluster memberships. (I) Box and whisker plot comparing overlap scores of data points from ESCHR hard clustering that were accurately assigned (orange) versus not accurately assigned (blue). The box shows the quartiles of the dataset, whiskers extend to 1.5*IQR, plotted points are outliers. Two-sided Mann-Whitney U test was used for statistical analysis. N = 126545, 750955 for inaccurate and accurate groups respectively. (J) Comparison of ESCHR overlap scores versus method agreement per each individual data point. X-axis is binned ESCHR overlap scores and y-axis is the average method agreement across all pairs of methods.

The second illustrative dataset, consisting of 15 gaussian distributions and 10,000 objects total, is prone to underclustering instead of overclustering. Similar to the first dataset, there is wide disagreement between the results of individual clustering algorithms using default parameters, but unlike the first dataset, all methods except SINCERA produced fewer than the actual number of ground truth clusters (Figs. 2e, S5b). As above, ESCHR overlap scores identify the regions of highest disagreement between the clustering methods (Fig. 2f), serving as a useful proxy for regions of uncertainty or difficulty in clustering, as quantified by F1 score (Fig. 2g). The soft clustering membership heatmap reveals overlapping relationships between clusters, identifying minimal overlap for ESCHR cluster 1, but a high degree of connectivity between ESCHR clusters 2, 3, 8, 10, and 7 (Fig. 2h).

In comparing the results for the 3-cluster and 15-cluster datasets, we observe that some clustering algorithms appear more prone to overclustering these synthetic datasets when used with their default hyperparameters, such as SINCERA and Scanpy, while other clustering algorithms are more prone to underclustering when used with their default hyperparameters, such as SC3. While these other clustering methods would undoubtedly produce higher accuracy results with manual hyperparameter tuning, ESCHR is able to produce high accuracy results without human intervention. Additionally, ESCHR provides a map of overlapping soft cluster assignments which gives insight into dataset structure and continuity, and the related overlap scores can be used to identify regions of uncertainty in cluster assignment due to ground truth cluster overlap.

We next sought to quantitatively evaluate the utility of ESCHR overlap scores across our full set of structurally diverse synthetic datasets with ground truth cluster labels. We first compared ESCHR overlap scores to the accuracy of assignment compared to ground truth labels per data point across all datasets and found that ESCHR overlap scores were significantly higher in inaccurately assigned cells (Fig. 2i). We then quantified the level of agreement between clustering assignments from all the different clustering algorithms we tested (Fig. 2a,e) and used this as an alternative external indicator against which to compare ESCHR overlap scores as a per-object metric for uncertainty and difficulty of clustering. This analysis revealed that higher ESCHR overlap scores corresponded to lower average method agreement per data point (Fig. 2j). Taken together, these initial comparisons demonstrate that ESCHR overlap scores identify meaningful uncertainty, and that when used in combination with the soft clustering results they enable more in-depth interpretation of dataset structure than other methods which produce only hard cluster assignments. Furthermore, ESCHR is able to provide these high-quality insights for datasets with diverse structural characteristics without the need for human intervention such as hyperparameter tuning.

### ESCHR outperforms other methods across measures of accuracy and robustness

We next performed systematic benchmarking of ESCHR against other clustering algorithms using a collection of 42 published real datasets in addition to the synthetic datasets described above. This collection of 42 published datasets vary widely in size (300-2,000,000 cells), source tissue (e.g. blood, bone marrow, brain), measurement type (sc/nRNA-seq, mass cytometry, flow cytometry, non-single-cell datasets), and data structure (varying degrees of discreteness and continuity) (Supplementary Table 3). For our evaluation criteria, we selected two extrinsic evaluation metrics, Adjusted Rand Index (ARI) and Adjusted Mutual Information (AMI), to assess two aspects of the clustering results: 1) accuracy and 2) robustness. Extrinsic evaluation metrics measure the distance of a clustering result to some external set of labels, and our two selected metrics ARI and AMI represent different approaches to this problem, with divergent biases. ARI tends to yield higher scores in cases of similarly sized clusters and similar numbers of clusters within and between the partitions being compared, while AMI is biased towards purity and yields higher scores when there are shared pure clusters between the two partitions (Methods) ^53^. Using ARI and AMI together should therefore provide a more complete comparison of clustering performance ^12,13^.

When we applied these extrinsic metrics ARI and AMI to assess clustering accuracy for our collection of synthetic datasets, ESCHR outperformed all other clustering algorithms across both metrics (Fig 3a). This superior performance was statistically significant for all cases except when compared to Seurat by ARI (Fig 3a, Supplementary Table 5). We also applied ARI and AMI to benchmark clustering accuracy in non-synthetic real datasets, although it is important to note that a priori known class labels do not generally exist for real-world single-cell datasets, and the various proxies accepted as ground truth labels should be interpreted with skepticism (discussed further in Supplementary Note 1). Keeping these caveats in mind, ESCHR still clustered real datasets more accurately by ARI than all methods except Agglomerative clustering, and the difference with Agglomerative clustering is not significant (Fig 3b, Supplementary Table 5). By AMI, ESCHR performance is significantly more accurate than SC3 and SINCERA and is on par with the remainder of the other methods tested (Fig 3b, Supplementary Table 5). Many of the ground truth labels that are widely accepted for real single-cell datasets are based on a hierarchical framework of clustering or manual labeling, which could explain why agglomerative clustering accuracy is higher relative to the other methods for the real datasets versus the synthetic datasets.

**Figure 3:**
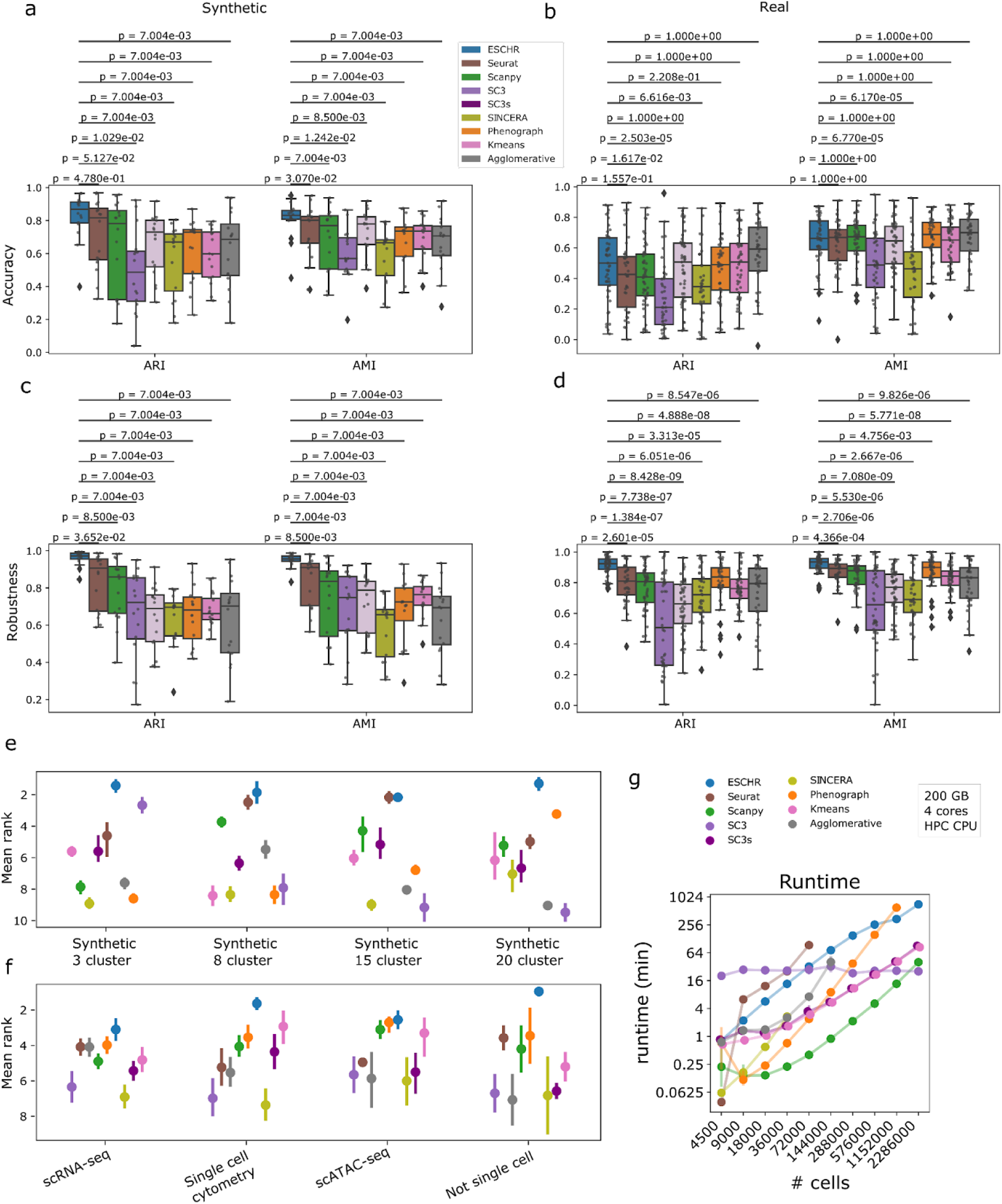
Systematic comparison of ESCHR clustering performance to competing methods in synthetic and real datasets. (A-D) Box and whisker plots comparing accuracy (A and B) and robustness (C and D) of results from ESCHR and all comparison methods across all synthetic (A and C) and real (B and D) benchmark datasets as measured by ARI (left) and AMI (right). Boxes show the quartiles of the dataset, whiskers extend to 1.5*IQR. Data points used in creation of box and whisker plots and shown in overlaid scatterplots are the means across 5 replicates for each dataset. Two-sided Wilcoxon signed-rank test with Bonferroni correction was used for statistical analysis comparing ESCHR to each method. N = 20 for comparisons using synthetic datasets and N = 42 for comparisons using real datasets. (E) Mean rank across all metrics shown in box-and-whisker plots for different categories of the synthetic datasets. Error bars show 1 standard deviation. (F) Mean rank across all metrics shown in box-and-whisker plots for different categories of the real datasets. Points represent means across all replicates of all datasets in a given category and error bars show 1 standard deviation. (G) Scalability comparison between ESCHR and other methods on synthetic datasets with increasing number of data points. X-axis is log scaled but labels show the unscaled values for easier interpretation. Each dot represents 5 replicates and error bars show 1 standard deviation. (Note that we experienced issues with our computing environment when running Seurat and we expect that Seurat should be capable of analyzing larger datasets within these memory constraints than what we were able to show here.)

After benchmarking for accuracy, we next used ARI and AMI to evaluate clustering robustness, by comparing results from repeated runs with different random subsamples of a given dataset (Methods). Due to its ensemble and consensus clustering approach, we expected ESCHR to perform well in these tests of robustness, and it demonstrated significantly better performance than all other clustering algorithms on both synthetic and real data across both ARI and AMI metrics (Fig. 3d,e, Supplementary Table 6). To gain insight into the generalizability of ESCHR versus the other methods for specific dataset types, we calculated the mean rank of each clustering algorithm across all metrics for major subcategories of our collection of datasets: scRNA-seq, flow/mass cytometry, multi-omic, and non-single-cell, as well as synthetic datasets of various structure. Different clustering algorithms perform better or worse for different subsets, but ESCHR is consistently ranked first or tied for first across these subcategories of both synthetic (Fig. 3e) and real datasets (Fig. 3f), indicating that its performance is more generalizable to diverse datasets than the other tested clustering algorithms.

We next evaluated the scalability of each method over a range of dataset sizes. While ESCHR generally takes the longest, this does not present a practical limitation for typical usage, as it is able to successfully complete analyses on millions of data points and the runtime scales linearly (Fig. 3g). This analysis also revealed that several of the alternative clustering algorithms we tested could not successfully run to completion for larger datasets. The dataset size limit for ESCHR is effectively the size limit of its underlying base clustering method, the Leiden algorithm implemented in Python ^46^. Furthermore, while it is not represented in most objective comparisons of method runtimes, it is important to consider that our method does not require the time-consuming process of manual hyperparameter exploration and selection. Therefore, in practical usage ESCHR may actually yield useful results more quickly than methods with faster runtimes. When taken together, these quantitative evaluations demonstrate that ESCHR performs favorably compared to the other methods tested here and achieves our desired goals of providing accurate and robust results, being generalizable to a broad range of diverse datasets, and being scalable to large datasets.

### ESCHR soft clustering and overlap scores reveal diverse structural characteristics of the MNIST dataset

To illustrate how ESCHR can identify regions of continuity and provide insight into cluster overlap and dataset structure, we selected the MNIST dataset for further analysis. This dataset, consisting of 70,000 handwritten digits with ground truth labels, is often used for machine learning demonstrations because the images can be visualized for intuitive interpretation ^35^. Other clustering algorithms set to default hyperparameters do not recapitulate the ground truth labels with high accuracy (Supplementary Fig. 6a), explained in part by the real variation that exists within the ground truth sets. For example, there are two common variations of the handwritten digit 1, and most of the clustering algorithms capture this difference. Of all the clustering algorithms tested, ESCHR clusters the MNIST dataset with the highest robustness and accuracy (Supplementary Fig. 6b), but it consistently splits the 1 and 9 digits into separate subsets (Fig. 4a,b), and in some cases it splits the digit 4 as well (Supplementary Fig. 6a). ESCHR usually produces highly consistent results from run to run thanks to its consensus clustering step, but this inconsistency around the digits 4 and 9 is suggestive of a high degree of continuity within and between these two classes (Supplementary Fig. 6c), which is highlighted by elevated ESCHR overlap scores in this region (Fig. 4c). The soft cluster membership heatmap also draws attention to the visual similarities between digits 3, 5, and 8, as well as the two types of handwritten 1 digits (Fig. 4d). These subset-level differences and connections between related digits motivated further investigation of the ESCHR outputs for the MNIST dataset.

**Figure 4:**
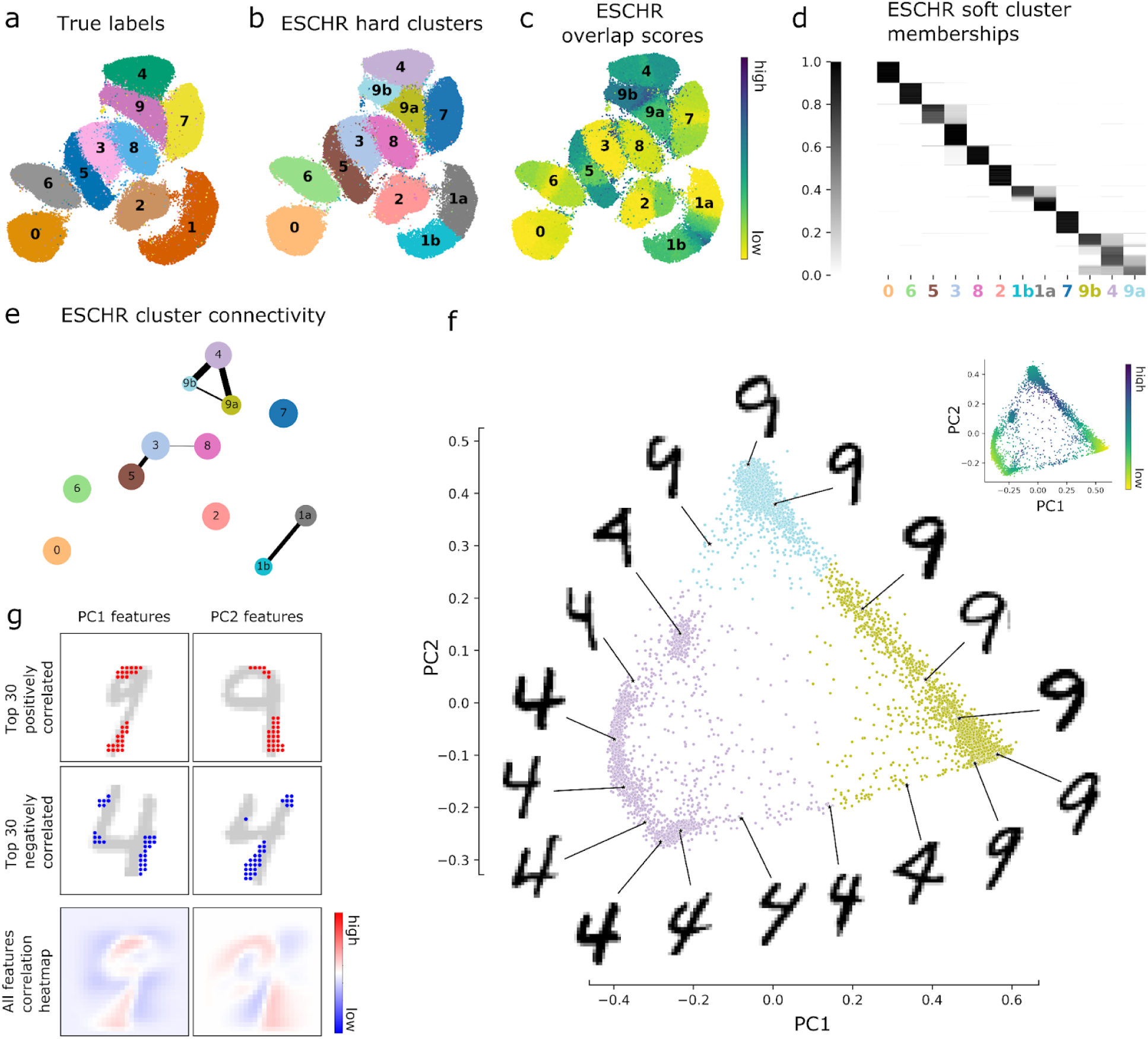
ESCHR performance on the classic benchmarking dataset MNIST. (A) UMAP visualization with points colored by true class labels. (B) UMAP visualization with points colored by ESCHR hard cluster labels. (C) UMAP visualization with points colored by ESCHR overlap score. (D) Heatmap visualization of ESCHR soft cluster memberships. (E) Nodes represent ESCHR hard clusters and are located on the centroid of the UMAP coordinates for all data points assigned to that hard cluster. Node size is scaled to the number of data points in a given cluster. Edges exist between nodes which were determined to have significant connectivity by ESCHR cluster connectivity analysis, and edge thickness is scaled to the connectivity score. (F) Visualization of data points from ESCHR clusters 4, 9a, and 9b projected onto the first two principal components resulting from PCA performed on the soft membership matrix of these three clusters. Primary scatterplot shows points colored by their ESCHR hard cluster assignment, and inset scatterplot shows points colored by ESCHR overlap score. Images are real examples from the MNIST dataset. (G) Scatterplot points in the first two rows of plots show the pixel locations of the 30 features with the largest positive (first row, red) and 30 largest negative (second row, blue) Pearson correlation to each of the PCs. Example digit images are underlaid in light gray to aid interpretation. The final row contains heatmaps with each pixel colored according to its Pearson correlation with PC1 (left) or PC2 (right), with bright red indicating a large positive correlation and dark blue indicating a large negative correlation.

To investigate the complex relationship between subsets of the digits 4 and 9 revealed by ESCHR, cluster connectivity mapping was applied to reveal significant overlap beyond what would be expected by random chance for the ESCHR clusters “4,” “9a,” and “9b” (Fig. 4e). Based on the soft membership heatmap, there appear to be some cells that are overlapping all three clusters, and some cells from clusters “9a” and “9b” that overlap separately with cluster “4” and not with each other (Fig. 4d). Unlike the simpler relationships between ESCHR clusters “3”-”5”-”8” and “1a”-”1b,” which could be analyzed by linear one-dimensional reduction, the more complex relationship around digits 4 and 9 could not be adequately captured or described along a single dimension, so principal components analysis (PCA) was applied to the ESCHR soft cluster memberships in order to reduce these relationships into two dimensions (Methods). Representative images selected from throughout the resulting PC space reveal that the between-cluster continuity is indeed reflecting the existence of a continuous progression through different conformations of the two digits 4 and 9 (Fig. 4f). Specifically, we can see that there is continuous progression through the 9’s based on how slanted they are, with two areas of higher density at either extreme. This explains why a clustering algorithm would be likely to split this into two clusters, albeit with a high amount of uncertainty about precisely where to make the split. The images also illustrate how the more slanted closed 4’s form a continuous transition primarily with cluster 9a and more vertically oriented closed 4’s form a continuous transition primarily with cluster 9b. This approach also allows us to identify features that are most correlated with the top two principal components. The top PC-correlated features lend further insight by identifying the specific pixels that are primarily capturing these changes in slantedness and upper loop closure (Fig. 4g). These analyses illustrate how structures within the MNIST dataset are not ideally suited for hard clustering assignment, but also how ESCHR is able to identify these structures and provide deeper insights than could be obtained by other hard clustering methods, or even beyond what is available from the ground truth class assignments.

### ESCHR captures cell types and continuity in static adult tissue

To illustrate how ESCHR can provide additional interpretability and insight for single-cell datasets, we selected an integrated scRNA-seq dataset of hypothalamic tanycytes ^39^ for further analysis. Tanycytes are elongated ependymoglial cells that form the ventricular layer of the third ventricle and median eminence, and have historically been classified into four subtypes (α1, α2, β1, β2) based on the hypothalamic nuclei where they project to, their spatial localization along the third ventricle, and their morphological, structural, genetic, and functional properties (Fig. 5a)^54^. More recent studies have suggested that many of these properties may exhibit substantial continuity between and within each of these subtypes ^36–40,55^. However, individual tanycyte scRNA-seq studies and an integrated analysis of these datasets all reported discrete groupings of tanycytes defined by hard clustering approaches ^37,39,56,57,58^, with no insight into the robustness of these assignments and whether there is overlap or continuity between them.

**Figure 5:**
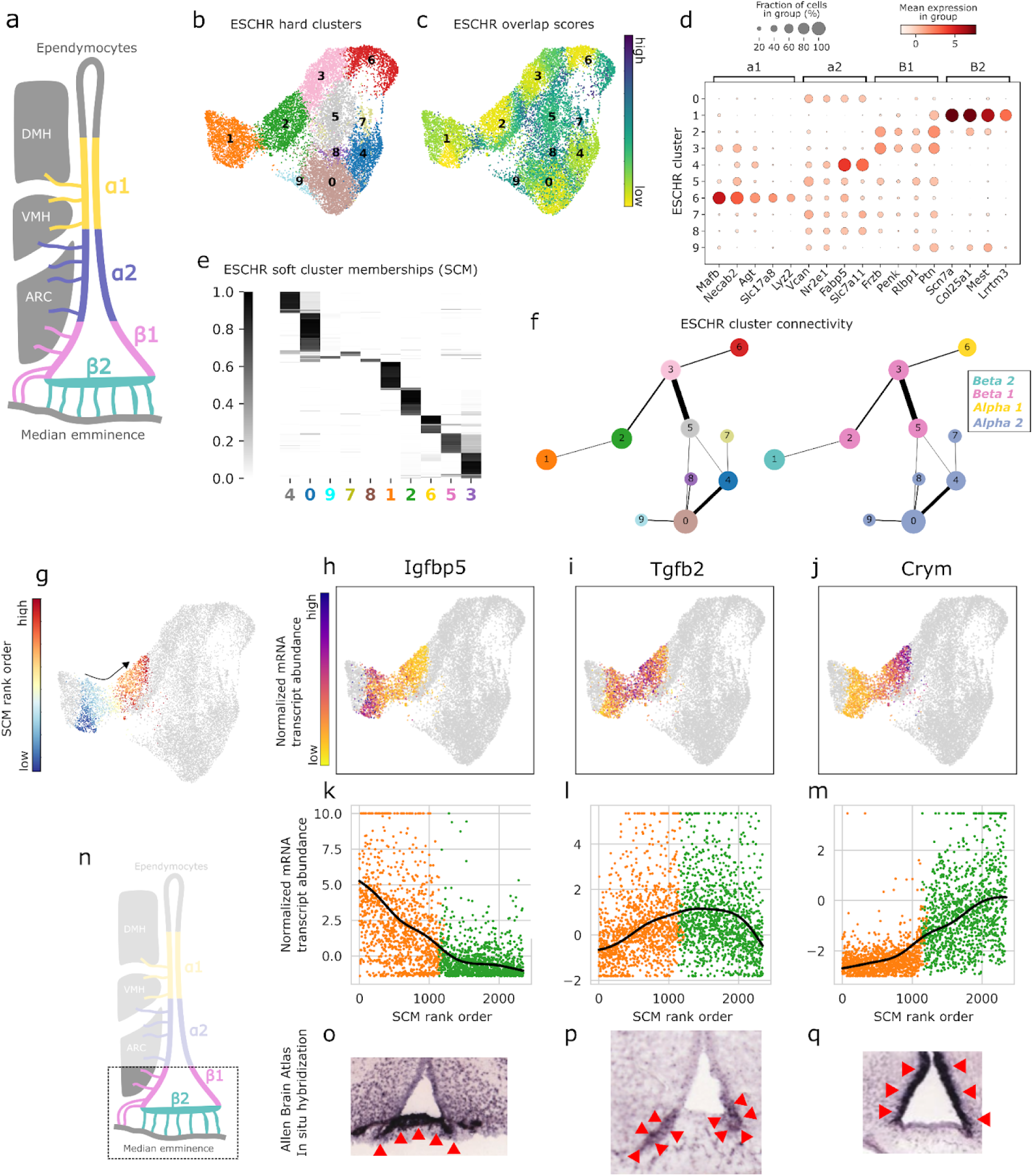
ESCHR describes continuity between and within canonical cell subtypes in static adult tissue. (A) Schematic illustration of canonical tanycyte subtypes in their anatomical context surrounding the third ventricle. (B) UMAP visualization with points colored by ESCHR hard cluster labels. (C) UMAP visualization with points colored by ESCHR overlap score. (D) Heatmap dotplot showing expression of marker genes for the canonical tanycyte subtypes across the ESCHR hard clusters. (E) Heatmap visualization of ESCHR soft cluster memberships (SCM). (F) Nodes represent ESCHR hard clusters and are located on the centroid of the UMAP coordinates for all data points assigned to that hard cluster. Node size is scaled to the number of data points in a given cluster. Edges exist between nodes which were determined to have significant connectivity by ESCHR cluster connectivity analysis, and edge thickness is scaled to the connectivity score. Node colors map to their ESCHR hard cluster colors from panel B (left) and to the color from panel A of the canonical subtype to which they primarily belong (right). (G) UMAP visualization with the subset of points which were included in the ordering analysis colored by ESCHR soft cluster membership (SCM) rank ordering score, and all others colored gray. (H-J) UMAP visualizations where points included in the ordering analysis are colored by their expression level and all others are colored gray. (K-M) Scatterplots showing normalized mRNA abundance on the y-axis and SCM rank order on the x-axis. Expression is bounded between the 2nd and 98th percentiles. Lines show gaussian-smoothed B-splines fit to the data. (N) Schematic illustration of the anatomical region being shown in O-Q. (O-Q) In situ hybridization (ISH) of coronal brain sections, using probes specific for Igfbp5, Tgfb2, and Crym (Allen Mouse Brain Atlas). Red arrowheads indicate the areas of expression in the region of interest.

Initial ESCHR analysis produced hard clustering outputs that match canonical tanycyte subtypes by their RNA expression profiles (Fig. 5b-d) ^39^. Subtypes β1 (expressing Fizb, Penk, Rlbp1, and Ptn) and α2 (expressing Vcan, Nr2e1, Fabp5, and Slc7a11) are represented by multiple hard clusters, while the subtypes β2 (expressing Scn7a, Cal25a1, Meat, and Lrrtm3) and α1 (expressing Mafb, Necab2, Agt, Slc17a8, and Lyz2) each correspond to a single hard cluster, indicating that there is more transcriptional diversity within the β1 and α2 populations. On top of this however, ESCHR overlap scores identify substantial heterogeneity within each hard cluster, including the β2 and α1 clusters (Fig. 5c), and the soft cluster memberships reveal additional levels of overlap and continuity between these canonical tanycyte subtypes (Fig. 5e). ESCHR cluster connectivity mapping (Methods) revealed significant overlap between the β1 clusters (2, 3, and 5) and each of the other three canonical subtypes (Fig. 5f). This result was somewhat unexpected, because transcriptional continuity was previously thought to exist only between spatially neighboring tanycyte subtypes ^38,55^. A more recent study provided evidence that β1 tanycytes exhibit some transcriptional continuity with both α1 and α2 tanycytes, but also indicated that β2 tanycytes were non-overlapping and transcriptionally distinct ^36^. Our analysis with ESCHR soft clustering memberships and cluster connectivity provide additional corroboratory evidence for the transcriptional continuity between β1 and α1/α2 tanycytes, but also reveal a previously uncharacterized relationship of transcriptional continuity between β1 and β2 tanycytes.

To further investigate this previously uncharacterized transcriptional overlap between β1 and β2 tanycytes, specifically between ESCHR clusters 1 and 2, we selected the subset of cells comprising the transitional zone between clusters, and rank ordered these based on whether their soft cluster membership was closer to β1 (ESCHR cluster 1) or β2 (ESCHR cluster 2) (Fig. 5g and Methods). Using this rank ordering scheme, we identified genes with expression patterns that correlate with progression through the transition zone from β2 to β1 tanycytes, either decreasing across the transition like Igfbp5 (Fig. 5h,k), peaking during the transition like Tgfb2 (Fig. 5i,l), or increasing across the transition like Crym (Fig. 5j,m). We next sought to determine whether these gene expression patterns in the transitional zone between ESCHR clusters were also observed in the spatial distribution of β2 and β1 tanycytes along the median eminence and third ventricle where these subtypes are thought to reside (Fig. 5n). To investigate this possibility, we examined the in situ hybridization (ISH) database from the Allen Mouse Brain Atlas (ABA; http://mouse.brain-map.org) ^59^ and observed that the overlapping expression for these three genes did in fact manifest as progressive spatial overlap spanning the anatomical regions canonically associated with β2 and β1 populations (Fig. 5o-q). Altogether, this analysis of tanycyte subtypes demonstrates the utility of ESCHR for 1) identifying robust and biologically meaningful hard cluster assignments, 2) providing insight into the overlap and continuity between cell type clusters, and 3) providing a springboard for further analysis of expression level transitions via soft cluster membership ordering.

## Discussion

Clustering is a fundamental tool for single-cell analysis, used to identify groupings of cell types or cell states that serve as the basis for direct comparisons between biological samples or between specific cell types within a biological sample, as well as numerous further downstream applications. However, it has proven challenging to generate appropriate and consistent cell groupings when using previously available clustering methods on single-cell datasets, due to 1) continuity and overlap between cell types, 2) randomness and stochasticity built into the clustering algorithms, and 3) non-generalizable hyperparameter settings that were optimized for a specific dataset or data type. To overcome these limitations we developed ESCHR, a user-friendly method for ensemble clustering that captures both discrete and continuous structures within a dataset and transparently communicates the level of uncertainty in cluster assignment. Using a large collection of datasets representing a variety of measurement techniques, tissues of origin, species of origin, and dataset sizes, we benchmarked ESCHR’s performance against several other clustering algorithms, demonstrating that ESCHR consistently provides the highest robustness and accuracy for clustering across all categories of this diverse dataset collection.

One of the key design features of ESCHR is a novel approach to hyperparameter randomization during the ensemble generation step. This eliminates the need for human intervention by the laborious process of manual hyperparameter tuning, which often relies on human intuition about which clustering results “look right” on a UMAP or tSNE plot during exploration of the clustering results by trial and error. The hyperparameter ranges for ESCHR were optimized empirically, and an important caveat to consider here is that exploring this parameter space was computationally expensive, limited by memory and runtime requirements, and there may be room for further optimization. Another direction not pursued here would be to generate specially customized hyperparameter ranges tailored to specific dataset categories, for example one set of hyperparameter ranges for scRNA-seq, and another set of hyperparameter ranges for mass cytometry, etc. One challenge for this strategy would be the lack of high quality ground truth labels for these specific dataset types (discussed further in Supplementary Note 1), and therefore one must be careful about what is being optimized for. Even in the best-case scenario of the MNIST dataset with high quality ground truth labels, our analysis revealed interpretable groupings and structure well beyond the ground truth labels (Fig. 4a-f), and optimizing ESCHR to only identify the ground truth labels here could have obscured these meaningful results. Overall, we believe our current more wide-ranging approach for setting the hyperparameter ranges is most appropriate, and the generalizability of ESCHR across a wide variety of datasets is a key advantage to the method.

A related area where we are more optimistic about potential improvement to the ESCHR algorithm would be expanding the number of hyperparameters randomized, in order to generate an even more diverse clustering ensemble. For example, we currently use k-nearest neighbor (kNN) graphs for the base Leiden clustering steps, but mutual nearest neighbor (mNN) or shared nearest neighbor (sNN) have shown good performance in other frameworks ^60^ ^61^ ^3^, and may improved ESCHR performance if incorporated as an additional hyperparameter to vary, or even as the sole method for graph generation in place of kNN. ESCHR may also benefit from expanding the set of distance metrics utilized. We currently restrict our analysis to euclidean and cosine distances due to their efficient implementations within our chosen fast approximate nearest neighbor (ANN) package ^62^. However, recent research has demonstrated the efficacy of a broader range of distance metrics for capturing diverse data structural properties ^63^. While not all of these metrics may be applicable in an ANN context, several may hold potential for enhancing the quality of our clustering outcomes. Finally, the current version of ESCHR uses only Leiden community detection for clustering in the ensemble stage, but additional base clustering methods could be explored and potentially incorporated in future versions.

Single-cell data is inherently complex and heterogeneous, and clustering methods often make assumptions about the structure of the data that may not hold in practice. For example, hard clustering methods assume discrete groups of single cells, which rarely exist in biological data ^15^. Many clustering algorithms make further assumptions about the shapes and other properties of these discrete groups. In the opposite direction, toward continuity rather than discreteness, numerous methods have been developed for trajectory inference in single-cell datasets ^4^, but these methods also make assumptions about dataset structure, for example many force a branched tree structure. ESCHR’s soft cluster outputs enable unified mapping of both discrete and continuous grouping structures, without the need for assumptions about the shape and properties of the dataset. To illustrate this concept, we used ESCHR to identify tanycyte subtypes and reveal the transitional continuity between them (Fig. 5a-q, Supplementary Fig. 7), which is notable because assumptions about lineage relationships or dynamic developmental processes in this static adult tissue would be inappropriate and could lead to inaccuracies and distortion. Instead, ESCHR can identify and characterize discrete and continuous patterns simultaneously, even in the same dataset, without relying on assumptions about data shape and properties.

One of ESCHR’s most useful outputs is the per-data-point overlap score, which enables users to estimate clustering uncertainty and interpret hard clustering results more effectively. This overlap score is derived from the degree of overlapping soft cluster assignments for each datapoint, which serves as a useful proxy for uncertainty and difficulty in cluster assignment. The Impossibility Theorem for clustering states that it is impossible for any clustering method to satisfy three proposed axioms of good clustering, and therefore all clustering algorithms must make trade-offs among the desirable features, and no clustering result can be perfect ^17^. Because of this, it is critical to evaluate the guaranteed uncertainty in a clustering result before using it for direct comparisons, downstream analyses, or hypothesis generation. ESCHR overlap scores provide a useful proxy for this uncertainty, which can be visualized alongside cluster assignments. We have validated the utility of these overlap scores by demonstrating that they are significantly higher for inaccurately assigned data points (Fig. 2k), and that they are positively correlated with the level of disagreement both between clustering algorithms (Fig. 2i) and between randomly subsampled replicates (Supplementary Figs. 7,8). Altogether, these findings demonstrate that ESCHR overlap scores provide meaningful insights into clustering uncertainty.

To make the advantages of ESCHR clustering easily accessible to the research community, we have made ESCHR available as a Python module on github (https://github.com/zunderlab/eschr), packaged as an extensible software framework that is compatible with the scverse suite of single-cell analysis tools ^31^. We have provided tutorials for how to incorporate it into existing single-cell analysis workflows as well as for how to use it as a standalone analysis framework. In conclusion, our results demonstrate that ESCHR is a useful method for single-cell analysis, offering robust and reproducible clustering results with the added benefits of per-cell overlap scores and soft clustering outputs for improved interpretability. By emphasizing ease of adoption, clustering robustness and accuracy, generalizability across a wide variety of datasets, and improved interpretability through soft clustering outputs and the quantification of uncertainty, we aim to support the responsible and informed use of clustering results in the single-cell research community.

## Methods

### ESCHR Framework

ESCHR takes as input a matrix, *M*, with *n* instances (e.g. cells) as rows and *p* features (e.g. genes/proteins) as columns. It does not perform internal normalization or correction, so input data are expected to have already been preprocessed appropriately. ESCHR can be thought of in three primary steps: ensemble clustering, consensus determination, and output/visualization.

Consistent with other published manuscripts in this domain, we will use the following notation. Let *X* = *x*_1_, *x*_2_,…, *x_n_* denote a set of objects, where each *x_i_* is a tuple of some *d*-dimensional feature space for all *i* = 1… *n*. Let *X* = *x*_1_, *x*_2_,…, *x_r_* denote a random subset of *X* where all of *x*_1_,…, *x_r_* are between 1 and *n*. ℙ = *P*_1_, *P*_2_,…, *P_m_* is a set of partitions, where each 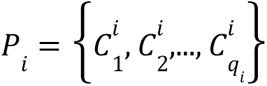 is a partition of an independent instantiation of *X_s_* and contains *q_i_* clusters. 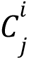 is the *j* th cluster of the *i* th partition, for all 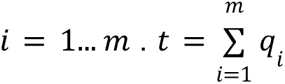 is the total number of clusters from all ensemble members. Where ℙ is the set of all possible partitions with the set of objects *X* and ℙ ⊂ ℙ_*X*_, the goal of clustering ensemble methods is to find a consensus partition *P*\* ɛ ℙ_*X*_ which best represents the properties of each partition in ℙ. Additionally, the more general terminology of “instance” and “feature” will generally be used rather than domain specific terms such as cells and genes/proteins.

### Hyperparameter-randomized ensemble clustering

The ESCHR ensemble is generated with Leiden community detection as the base clustering algorithm ^46^. Leiden is applied using Reichardt and Bornholdt’s Potts model with a configuration null model ^64^. Diversity is generated amongst ensemble members through a combination of data subsampling and Leiden hyperparameter randomization. The subsampling percentage varies for each ensemble member and is selected from a gaussian distribution with the mean μ scaled to dataset size within the range [30,90]. After subsampling a random subset *X_s_* from *X*, principal components analysis (PCA) is applied to generate the most informative features for this data subspace. For sparse datasets, truncated singular value decomposition (no mean centering) is applied in place of PCA. In the subsequent clustering step, three numerical hyperparameters are randomized for each ensemble member: 1) *k*, the number of neighbors for building a k-nearest neighbors (kNN) graph, 2) the choice of distance metric for building the kNN graph, and 3) *r*, a resolution parameter for the modularity optimization function used in Leiden community detection. The numerical hyperparameters *k* and *r* are randomly selected from within empirically established ranges [Supplementary Fig. 2]. The distance metric is selected between either euclidean or cosine, because these choices are efficiently implemented for fast calculation of approximate nearest neighbors (ANN) in our chosen implementation, nmslib ^62^. Since each ensemble member is independent, we implemented parallelization via multiprocessing for this stage of the algorithm. Ensemble size is set at a default of 150 based on experiments demonstrating that this was sufficient to reach convergence to a stable solution for all of our diverse collection of datasets [Supplementary Fig. 2].

#### Bipartite graph clustering and consensus determination

Bipartite graph clustering was used to obtain consensus clusters from the ESCHR ensemble. This approach was selected because methods that compute consensus using unipartite projection graphs of either instance or cluster pairwise relations suffer from information loss ^48^. For these calculations, the biadjacency matrix is defined as: 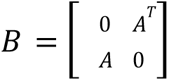 where *A* is an *n* × *t* connectivity matrix whose rows correspond to instances {1 … n} and columns correspond to the ensemble clusters {1 … t}. *A_i,j_* is an indicator that takes value 1 if instance *i* belongs to the *i*-th cluster and 0 otherwise. Using this, we then create a bipartite graph *G* = (*V*, *W*). The weights matrix *W* = *B*, and *V* = *V*_1_ ∪ *V*_2_, where *V*_1_ contains *n* vertices each representing an instance of the data set *X*; *V*_2_ contains *t* vertices each representing a cluster of the ensemble. Given our bipartite graph *G*, we can define a community structure on *G* as a partition 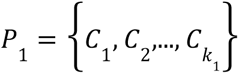 containing pairwise disjoint subsets of *V*_1_ and 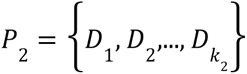 containing pairwise disjoint subsets of *V*_2_, such that all *V_1_* nodes in a specific *C_i_* are more connected to a particular subset of *V_2_* than the rest of the nodes in *V_1_* are, and likewise (but opposite) for a given *D_j_* of *V_2_*. Optimal *P*_1_ and *P*_2_ are computed with the Leiden algorithm for bipartite community detection with the Constant Potts Model quality function ^65^ ^47^. This approach was designed to overcome known issues with previous bipartite community detection approaches, including the resolution limit and the assumption that the optimal partitioning will have an equal number of communities for each node type ^66^ ^67^. There is one tunable hyperparameter for this approach, the resolution γ. To avoid the need for external hyperparameter tuning, we implement an internal hyperparameter selection strategy. Specifically, we perform clustering for a range of γ values and select the result which maximizes the sum of ARI values to all other clusterings to return as the primary result from consensus clustering. To obtain the final consensus result, we collapse the base ensemble clusters contained in *V*_2_ into the *P*_2_ meta-cluster to which they were assigned. This results in each vertex of *V*_1_ having a weighted edge to each meta-cluster equal to the sum of its edges with constituent base clusters of *V*_2_. The resulting weighted bipartite graph *G*\* therefore represents the final consensus clustering *P*^∗^, with *n* vertices representing the instances, *q*\* vertices representing the final consensus clusters, and weighted edges representing the membership of instance *i* in each of the *q*\* clusters of *P*^∗^.

#### Hard and soft clustering outputs

Let Θ ɛ ℝ *n* × *q*\* be a nonnegative matrix where each row, 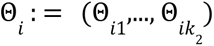, contains nonnegative numbers that sum to less than or equal to one, representing the membership of instance *i* in each of the *q* clusters of *P*. Θ_*ij*_ is calculated by dividing the weight of the edge between instance *i* and consensus cluster *D_j_* by the sum of all edge weights for instance *i*. We refer to this matrix as the soft membership matrix and to each row as the association vector *v* for each instance. To determine hard clustering assignments, each instance is assigned to the meta-cluster with the highest entry in its association vector *v*, with ties broken randomly. A “core” cell of a given cluster *j* will have Θ*_ij_* = 1 and zeros elsewhere, while a “transitional” instance may have up to *q\** non-zero membership values. To describe the degree to which a given instance is “core” versus” transitional”, we define an “Overlap Score”, Ω, for each instance as the highest membership value in its association vector ( Ω = *max*(*v*)). We can additionally calculate the mean of all instance memberships in a given cluster to yield a measure of each cluster’s discreteness, which we call the “cluster stability score” 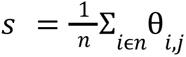.

### Cluster connectivity mapping

To map the connectivity structure of clusters, we first calculate the sum-of-squares-and-cross-products matrix (SSCP) of the soft membership matrix Θ, which is calculated as *S* = Θ’Θ and then consider *S*_*i*,*i*_ to be an uncorrected measure for connectivity between consensus clusters *i* and *j*. To correct for connectivity that may result from random chance, we first estimate a null distribution of connectivity scores accounting for the following attributes of Θ: (1) the association vector *v* for a given instance are proportions and can sum to no more than 1 (with cells summing to less than one being potentially outliers) and (2) the distribution of values is not uniformly distributed and will be differently skewed for different datasets depending on overall levels of continuity or discreteness. In practice, we achieve this by independently shuffling the association vector, *v*, for each instance to generate a random sample. We then calculate the SSCP for 500 iterations of this randomization procedure. Using this empirical null distribution, we then calculate a p-value for each observed edge and prune edges that do not meet a default alpha value cutoff of 0.05. Thus the final corrected connectivity is defined as the ratio of the cross-product of instance memberships between a given two clusters normalized to the cross-product of instance memberships expected under constrained randomization.

### Exploring soft cluster continuity for the Tany-Seq dataset

To visualize transitional zones between connected clusters in the Tany-Seq dataset ^39^, we devised a simple approach for creating a one-dimensional ordering of the instances in a transitional zone based on their membership in the connected clusters of interest. Specifically, the ordering score, δ*_*i*_*, of cell *i* having *m_j_* membership in the cluster at position *i* along the cluster path of interest was calculated as: 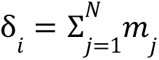, where *N* is the number of clusters in the path. To obtain the relevant cells of interest for ordering, we used the following criteria: cells were included if they had (1) >90% membership in either one of the 2 clusters of interest or (2) >5% membership in both clusters and a combined membership of >80% in the 2 clusters. To identify marker features associated with the one-dimensional soft cluster transition paths in Figure 4, we calculated the Pearson correlation between each feature and the vector of cluster memberships for each cluster in the path. Features were then selected based on their correlation with each of the clusters individually and based on the sum of their correlations across the clusters. The three genes in Figure 4 H-J were selected from the top ten features identified through each of these methods based on their expression patterns and the availability of in situ hybridization images of sufficient quality in the Allen Mouse Brain Atlas ^59^. To handle outliers for the expression heatmap UMAP plots and the expression scatterplots in Figure 4 H-J, values were thresholded to fall between the 2nd percentile and 98th percentile. The curves overlaid on the expression scatter plots in these panels were generated by first fitting B-splines with degree 3 (cubic) to the points included in the scatterplot. To generate a smoothed curve, a gaussian kernel with sigma of 10 was applied on the results of the spline function evaluated at 100 evenly spaced points within the range of the number of points included in the scatter plot. This is approximately equivalent to the behavior for large data sizes of the ′geom_smooth′ function from the R package ggplot ^68^.

### Exploring soft cluster continuity for the MNIST dataset

While a linear ordering approach could in principle be used to create an ordering across a path of more than two connected clusters, it would likely only be effective in cases where connectivity mapping identifies a linear path of successively connected clusters. In cases like the example in Figure 4 with the MNIST dataset ^35^, where connectivity mapping identified a ring of 3 connected clusters, it will generally not work as well since a group of more than two clusters with nonlinear connectivity may exhibit more complex continuity structures than could be captured with a simple linear ordering. We therefore devised another method for distilling the core continuity structure for cases of greater than two clusters and nonlinear connectivity paths. We first performed principal components analysis (PCA) on the columns of the soft membership matrix Θ that correspond to the hard clusters selected for analysis, thereby capturing the primary axes of variation contained within these soft memberships. We then projected the data onto the first two PCs and used this to gain insight into the continuity structure by (1) visualizing the data points belonging to the relevant hard clusters projected into the space of these first two PCs and (2) identifying and exploring the features most highly correlated with these PCs.

### Clustering evaluation metrics

Extrinsic evaluation metrics measure the distance of a clustering result to some external set of labels. When these labels are ground truth class labels, we can consider these to be measures of accuracy. However, they can also be used in other contexts such as with an “external” set of clustering labels. There are numerous metrics that can be used to measure this distance between a given set of predicted cluster labels and a ground truth or other set of external labels. Each of these metrics introduces some type of bias in evaluating the accuracy and robustness of clustering results, as discussed further below. To diversify these biases, we selected metrics from each of the three different categories of existing metrics: F-measure from the suite of metrics calculated by matching sets, Adjusted Rand Index (ARI) from the methods that employ peer-to-peer correlation, and Adjusted Mutual Information (AMI) from the information theoretic measures ^53^. We use the F1-score to evaluate per-true cluster accuracy for the example synthetic datasets in Figure 2, and we use ARI and AMI to evaluate accuracy and robustness in our systematic benchmarking in Figure 3 and in Supplementary Figures 2 and 6.

The F-measure is used to measure per-class accuracy and balances the importance of false negatives and false positives by taking the harmonic mean of precision and recall. When they are balanced with a ratio of 1, this is called the F1-score, and this is the version used in this manuscript. To apply this in the context of evaluating accuracy for a given clustering per each ground truth cluster, we must first determine the best matching predicted cluster for each ground truth cluster, which is discussed further below. We then calculate the F1-score per ground truth cluster and best-matched predicted cluster as follows:

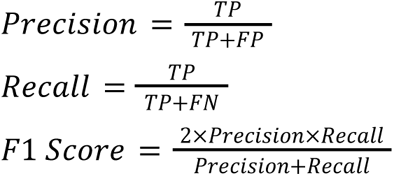

where *TP* is the number of data points overlapping between the ground truth cluster and predicted cluster, *FP* is the number of data points in the predicted cluster and not the ground truth cluster, *TN* is the number of data points in neither the predicted cluster nor the ground truth cluster, and *FN* is the number of data points in the ground truth cluster and not the predicted cluster. We use the Hungarian algorithm to search for the best mapping of predicted clusters to ground-truth clusters ^69^. Label matching when there are different numbers of labels is an unsolved problem and is the primary source of bias for this metric. This metric is not adjusted for chance, which is another potential source of bias. It is for these reasons that we only used this in the case where we required a measure of accuracy per true cluster and did not also apply this metric in our systematic benchmarking to compare quality of results between different methods. The values calculated by this measure can range from 0 to 1, with 1 indicating perfect accuracy. F1 scores were calculated using the implementation in sklearn (v 1.0.1).

The ARI is the corrected-for-chance version of the Rand index, which measures the agreement between two sets of partition labels *U* and *V* ^70^. The ARI is defined as:

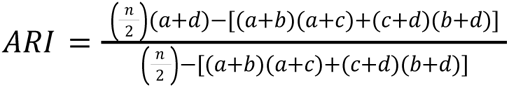

where *a* is the number of pairs of two objects in the same group in both *U* and *V*; *b* is the number of pairs of two objects in different groups in both *U* and *V*; *c* is the number of pairs of two objects in the same group in *U* but in different groups in *V*; and *d* is the number of pairs of two objects in different groups in *U* but in the same group in *V*. Random clusterings have an expected score of zero and identical partitions have a score of 1. ARI is biased towards solutions containing (1) balanced clusters (i.e. similar size clusters within each partition) and (2) similar cluster numbers and sizes between the two partitions ^13^. ARI was calculated using the implementation in sklearn (v 1.0.1).

AMI is the corrected-for-chance version of Mutual Information, which quantifies the amount of information that can be obtained about one random variable (in this application, a list of cluster labels) by observing the other random variable (another list of cluster labels) ^71^. Let *C* = {*C*_1_, *C*_2_,…, *C_tc_*} and *G* = {*G*_1_, *G*_2_,…, *G_tg_*} be the predicted and ground truth labels on a dataset with n cells. AMI is then defined as:

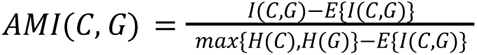

Here *I*(*C*, *G*) represents the mutual information between *C* and *G* and is defined as: 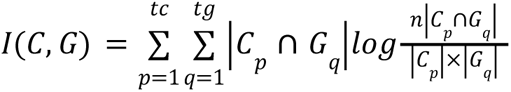. *H*(*C*) and *H*(*G*) are the entropies: 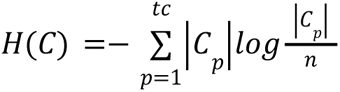 and 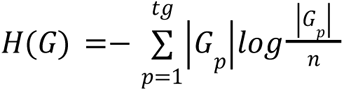. *E*{*I*(*C*, *G*)} is the expected mutual information between two random clusters. Random clusterings have an expected score of zero and identical partitions have a score of 1. AMI is biased towards solutions containing pure clusters, with a “pure cluster” being defined as a cluster in one set of labels that contains instances from only one cluster of the other set of labels to which it is being compared ^13^. AMI was calculated using the implementation in sklearn (v 1.0.1).

### Systematic Benchmarking

For benchmarking ESCHR, we selected the following widely used clustering algorithms for comparison: (1) SC3 (version 1.10.1 from Bioconductor) ^24^, (2) Seurat (version 4.1.1 from CRAN) ^72^, (3) SINCERA (script adapted from https://www.singlecellcourse.org/biological-analysis.html) ^73^, (4) Scanpy (version 1.8.2 from Anaconda) ^33^, (5) Phenograph (version 1.5.7) ^60^, (6) SC3s (version 0.1.1 through Scanpy) ^25^, and (7&8) K-means and agglomerative hierarchical clustering (from scikit-learn version 1.0.1) ^74^. Clustering algorithms were excluded from our benchmarking comparison if they did not meet the following selection criteria: (1) software freely available; (2) code publicly available; (3) can run on multiple data modalities (e.g. not scRNA-seq-specific); (4) no unresolved errors during install or implementation; (5) does not require additional user input during the algorithm (other than prior information); and (6) able to complete analysis of datasets with >= 100,000 data points and 2,000 features.

For included methods, we followed the instructions and tutorials provided by the authors of each software package. For K-means, SC3s, Agglomerative methods which require pre-specification of cluster number, we calculated distortion scores over a range of cluster numbers for K-means clustering and used the elbow method to select the optimal cluster number for use across all three methods. Default values were used for all other hyperparameters for each tool, as is common practice for most realistic use cases ^6^ ^75^. ESCHR was also run with all default settings. For all benchmarking analyses, the memory was set to 200GB of RAM on the University of Virginia (UVA) Rivanna High Performance Computing (HPC) cluster.

No random seeds were intentionally fixed, but from inspecting the respective codebases we believe it is likely that there remained internally-fixed random seeds for some functions within some of the tested methods. Many common methods have internally-fixed random seeds and/or default hyperparameters with fixed random seeds. This practice may mask a lack of robustness of these methods, and should only legitimately serve to replicate exact analyses when that is desired by the end user.

To assess accuracy of methods in clustering our synthetic and image datasets, which have ground truth labels, we used the two extrinsic evaluation metrics defined above (ARI, AMI). For these purposes, each of five independent runs of a given method was scored against the ground truth labels. Since it is nearly universal in papers describing new single-cell analysis methods, we also applied this analysis to evaluate accuracy for our collection of “real” datasets using available published labels. However we stress that we do not think this is a reliable or effective measure for evaluating new clustering methods, as we detail further in Supplementary Note 1.

We also used the extrinsic metrics ARI and AMI to evaluate the stability and reproducibility of hard clustering results. In line with standard practice for benchmarking stability/robustness ^76^, we performed repeated runs with 5 random subsamples (90%) of each dataset for every method. This simulates slight differences in data collection and/or preprocessing, and if the clustering is capturing a true underlying structure rather than overfitting to noise it should be detected regardless of the exact set of cells that are sampled for the analysis. We then calculated pairwise scores for each metric between each of the 5 independent runs of a respective method and then took the mean across replicate pairs to obtain the final score per dataset-method.

To calculate both method and replicate “agreement scores” for comparison with ESCHR overlap scores (Fig. 2j), we first constructed contingency matrices between all pairs of replicates and methods and mapped the cluster labels from the result with more clusters to the result with fewer clusters. Using the shared labels between a given pair of clustering results we could then calculate per-instance agreement (binary) within the pair of results. The final per-instance score was calculated as the mean agreement across all possible combinations.

We similarly calculated the F-measure per true cluster per method (Figure 2 c,g) by constructing a contingency matrix between the true cluster labels and the hard cluster labels predicted by each method. Using this mapping we then calculated the harmonic mean of precision and recall separately for each true cluster and its best-matched predicted cluster.

### Statistical analyses

Statistical comparisons were performed using the “scipy.stats” and “statannotations” python packages ^77^ ^78^. The two-sided Wilcoxon signed-rank test with Bonferroni correction was used to compare the performance of ESCHR versus each alternative method in the systematic benchmarking panels shown in Figure 3. Comparisons were calculated using dataset means across replicates for all tested datasets. N = 20 for comparisons using synthetic datasets and N = 42 for comparisons using real datasets. The two-sided Mann-Whitney-Wilcoxon was used for comparing overlap scores between accurately and inaccurately assigned cells in Figure 2I, with N = 126545 and N = 750955 for inaccurate and accurate groups respectively. The resulting p-value was below the threshold of calculation in a standard python computing environment and was reported as zero, so we have reported this as p<0.00001 in the figure.

## Data availability

All “real” datasets are publicly available. The 42 datasets used in our study are described in Supplementary Table 2, including information and links to download preprocessed datasets. The only processing step we performed was log2 transformation for scRNA-seq datasets and arcsinH transformation for mass cytometry datasets if a given dataset was not yet scaled. The MNIST dataset was downloaded from keras datasets and was preprocessed as recommended by the accompanying documentation.

Synthetic datasets were generated using [sklearn “make_blobs”](https://scikit-learn.org/stable/modules/generated/sklearn.datasets.make_blobs.html) using various combinations of object number, feature number, cluster number, cluster size, cluster standard deviations, and cluster anisotropic transformations. These datasets are available at: https://doi.org/10.5281/zenodo.10261863

ISH images were downloaded from the ABA portal (https://mouse.brain-map.org/) and are freely available.

Further information and requests for resources should be directed to and will be fulfilled by Eli Zunder.

## Code availability

The ESCHR python package can be downloaded from PyPi (TBD), github (https://github.com/zunderlab/eschr). The version used for making the figures in this manuscript is noted in the documentation. ESCHR is compatible with standard python single-cell data structures and can be easily incorporated into scverse workflows or used as a standalone framework.

## Acknowledgments

Research reported in this publication was supported by the National Institute of Neurological Disorders and Stroke of the National Institutes of Health under award number R01NS111220 to E.R.Z. The content is solely the responsibility of the authors and does not necessarily represent the official views of the National Institutes of Health. S.M.G. was further supported by the Transdisciplinary Big Data Science Training Grant, award number T32LM012416. The authors acknowledge Research Computing at The University of Virginia for providing computational resources and technical support that have contributed to the results reported within this publication. URL: https://rc.virginia.edu. The authors would also like to thank Sean Chadwick for critical feedback on the manuscript.

## Author Contributions

S.M.G and E.R.Z. conceptualized the ESCHR clustering method. S.M.G. developed the ESCHR Python package. S.M.G. and E.R.Z. planned all analysis and benchmarking experiments. S.M.G. performed all analysis and benchmarking experiments, and prepared figures. S.M.G. and E.R.Z. wrote the manuscript.

## Declaration of Interests

The authors declare no competing interests.

## Notes

### Competing Interest Statement

The authors have declared no competing interest.

