## Supplemental Information for "A hyperparameter-randomized ensemble approach for robust clustering across diverse datasets"

### Supplementary Notes

#### 1. On the difficulty of identifying a "ground truth" to assess clustering accuracy on single-cell datasets

Clustering accuracy can be evaluated with extrinsic validation metrics to compare a set of cluster labels to a set of ground truth class labels. This requires the existence of a priori known ground truth class labels, which generally do not exist for real-world single-cell datasets. Nevertheless, clustering accuracy is widely used as a key metric for evaluating single-cell clustering methods, and many publications featuring new single-cell clustering methods have claimed to measure the accuracy of their clustering results using a variety of different annotations as proxies for ground truth. These "ground truth" annotations are frequently based on either manual annotation and/or previous clustering results, and we are skeptical about the validity of these human-guided ground truth assignments. There are some rare exceptions where ground truth class labels in single-cell data are unambiguous and clear cut, such as unique cell lines or drug treatments. But in most cases, we propose that the ground truth annotations used to benchmark clustering accuracy in single-cell datasets are inappropriate, and not a useful yardstick to measure the "goodness" of clustering.

Besides cell lines and drug treatments, all other examples of "ground truth" annotations of single-cell data that we're aware of are problematic. Previous clustering results, even those obtained with human-optimized parameter tuning to give a result that looks best the human eye on a tSNE or UMAP plot, are still just one result from a single clustering algorithm, and should not be the yardstick by which all other clustering algorithms are judged. Even expert-guided "ground truth" annotations on a cell-by-cell basis still rely on our imperfect knowledge of functional markers, and rely on human intuition which doesn't perform well in high-dimensional space. An important additional point which has been made by others is that manual annotation is inherently limited to only capturing already-known biology and will miss novel findings [1] [2].

Another annotation that has previously been used as ground truth is patient source of tissue samples, but if these samples include a heterogenous mixture of cells then it is not reasonable to expect that the ground truth that clustering should capture would be differences between patients rather than between cell types shared by all patients. Developmental stage or collection day of differentiating cell types is also commonly considered to be a "gold standard" annotation, but considering that cell types often persist over varying time spans of development or other dynamic biological processes, it is a flawed assumption to believe that clustering should ideally be capturing differences between stages or collection days. Other studies use some versions of "sorting" by an alternate data modality as true labels. A common example of this is using flow cytometry-based sorting using a small set of cell surface proteins to select out different populations of cells and then using these population labels as ground truth for assessing clustering of scRNA-seq of the cells. While this can certainly be useful for a researcher aiming to explore mRNA expression specifically within cells defined by protein markers, there is no reason to believe that the actual true best clustering based on one modality should capture the same population structure that is defined by another modality, and this is therefore also not a valid basis for assessing the accuracy of a clustering result in the context of evaluating a new method.

#### Supplementary Figures

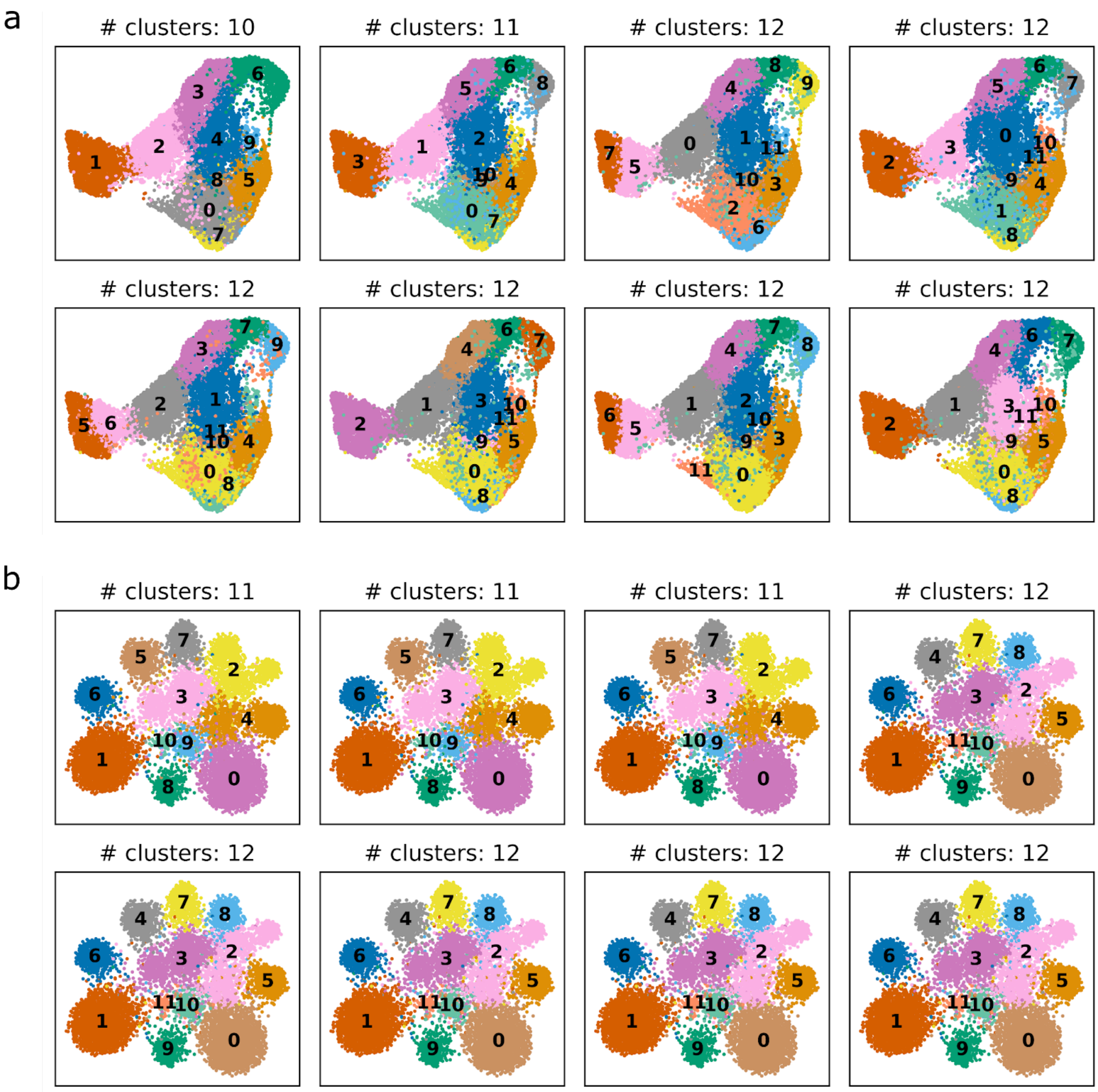

**Supplementary Figure 1: Inherent randomness in the commonly used Leiden algorithm.** UMAP visualizations of 8 replicates of Leiden clustering run with identical hyperparameters on identical input k-NN graphs. (A) shows the Tany-seq scRNA-seq dataset [3] which was explored in Figure 4 and (B) shows this analysis applied to the synthetic gaussian 10 cluster dataset that was used as one of the examples in Figure 2. Points are colored by cluster labels and numerical cluster labels are placed on the centroid of the cluster.

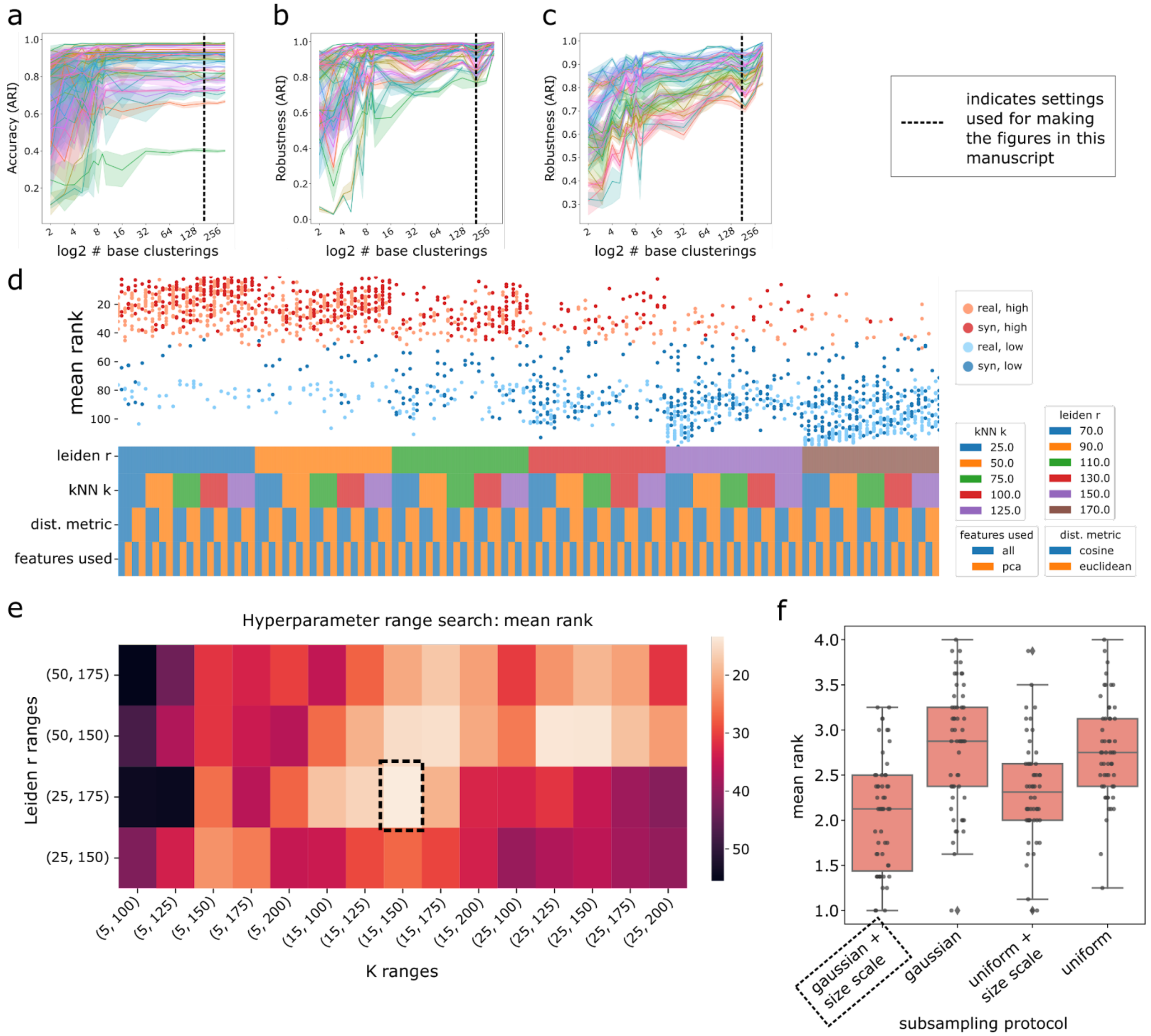

**Supplementary Figure 2: Establishing default settings for ensemble hyperparameter randomization.** (A-C) Line plots showing quality metrics on the y-axis and ensemble size on the x-axis, with synthetic datasets evaluated on accuracy by ARI (A) and robustness by ARI (B) and real datasets evaluated on robustness by ARI (C). Each dataset is represented by a different color of line. Dark lines show the mean and lighter colored surrounding band shows 1 standard deviation from across 5 replicates of ESCHR hard clustering results. X-axis is log2 scaled but the text labels show the unscaled values to aid interpretation. (D) Composite visualization illustrating the variability of optimal hyperparameters across different datasets. Dark and light red dots represent the highest scoring 10 hyperparameter combinations for each synthetic dataset and real dataset respectively. Dark and light blue dots represent the lowest scoring 10 hyperparameter combinations for each synthetic dataset and real dataset respectively. The vertical position of the dots represents the mean rank across accuracy ARI and AMI and robustness ARI and AMI and the horizontal position indicates the combination of hyperparameters represented by a given point, as specified by the overlapping colorbars. (E)

Heatmap showing the mean rank across accuracy ARI and AMI and robustness ARI and AMI and across all synthetic datasets analyzed with different ranges set for the 2 numeric hyperparameters, k number of neighbors for building the k-NN graph on the x-axis and r resolution for Leiden clustering on the y-axis. Dashed box indicates the highest scoring combination, which was selected for use in the version of ESCHR that was used to generate the figures in this manuscript. (F) Box and whisker plot showing the mean rank calculated as described for panels D and E for each of 4 different subsampling protocols. Individual dots represent each of 5 replicates of each of the 20 synthetic datasets. Note that lower rank indicates better performance.

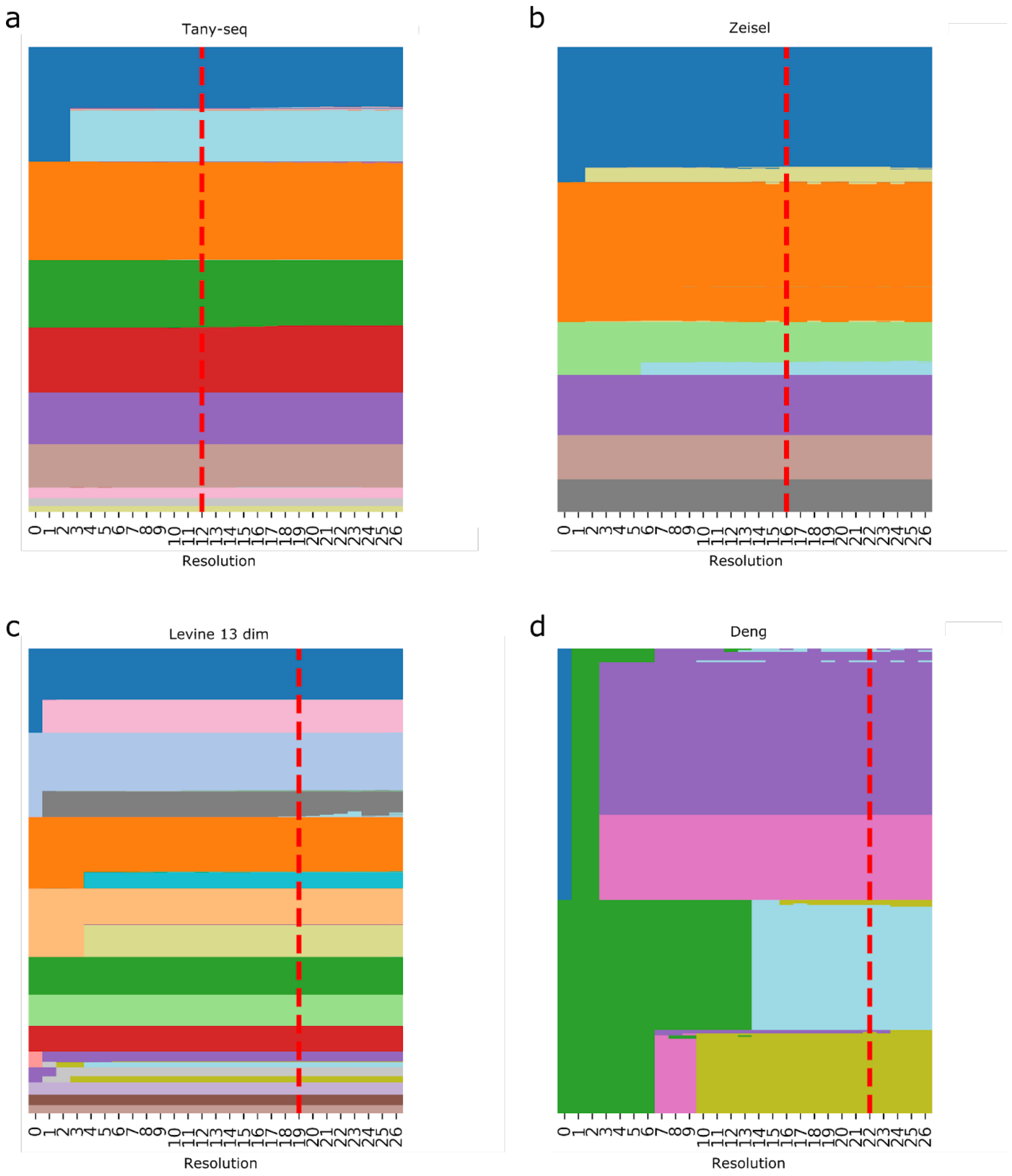

**Supplementary Figure 3: Validation of internal hyperparameter selection at consensus stage.** Sorted heatmap visualizations indicating ESCHR hard cluster groupings (y-axis, sorted group labels together) across different resolutions (on x-axis) for the consensus stage bipartite clustering. Example datasets shown here include the Tany-seq scRNA-seq dataset [3] (A), Zeisel scRNA-seq dataset [4] (B), Levine 13 dim mass cytometry dataset [5] (C), and the Deng scRNA-seq dataset [6] (D). The red dashed line indicates the resolution that was selected by the ESCHR internal optimization protocol in each case.

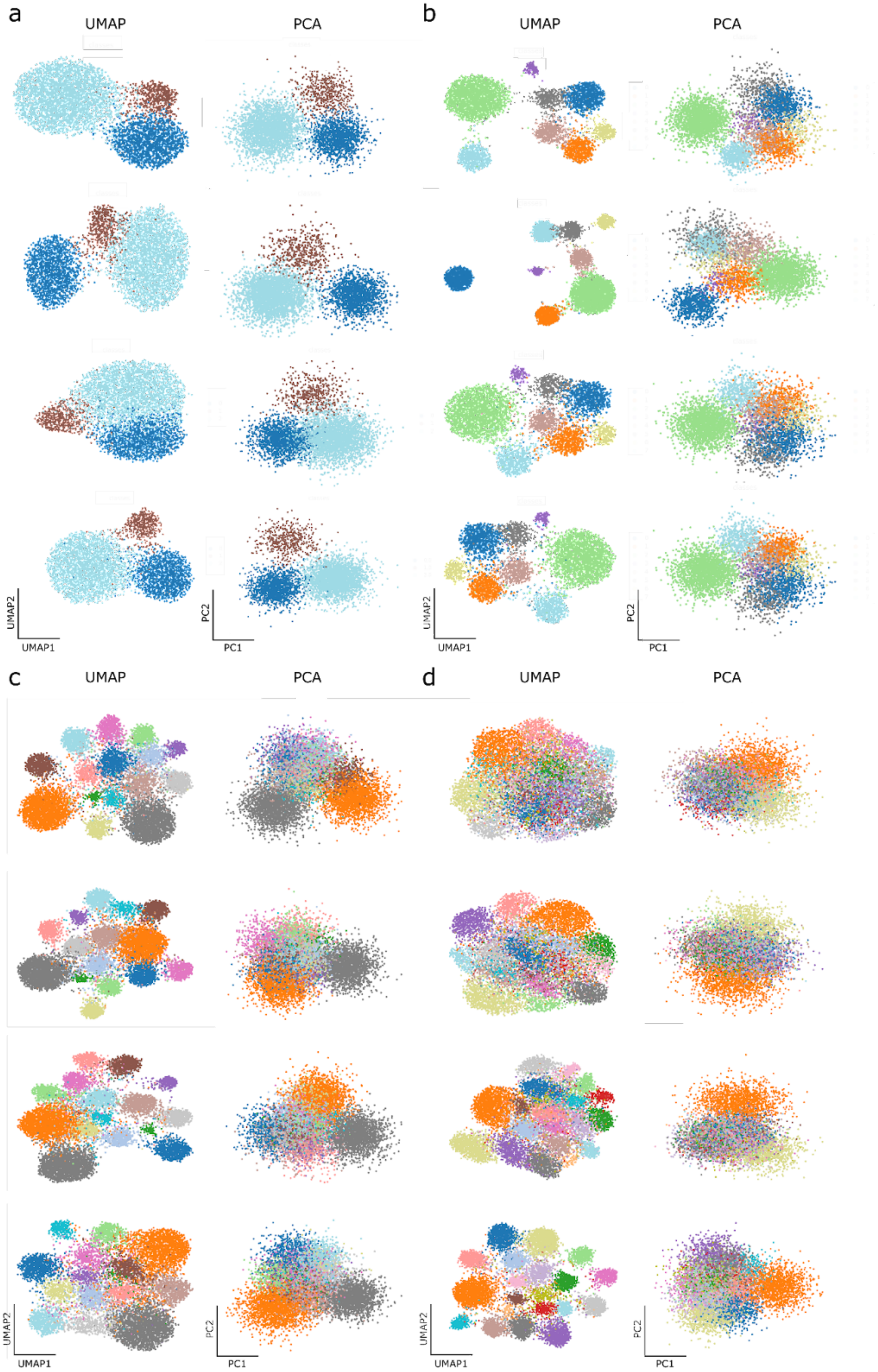
a
UMAP
PCA
b
UMAP
PCA
UMAP2
UMAP1
PC2
PC1
UMAP2
UMAP1
PC2
PC1
c
UMAP
PCA
d
UMAP
PCA
UMAP2
UMAP1
PC2
PC1
UMAP2
UMAP1
PC2
PC1

**Supplementary Figure 4: 2D visualizations of synthetic datasets.** UMAP (first column) and PCA (second column) dimensionality reduced 2D visualization of the synthetic gaussian datasets used in Figures 2 and 3. (A) contains the datasets with three ground truth clusters, (B) contains the datasets with 8 ground truth clusters, (C) contains the datasets with 15 ground truth clusters, (D) contains the datasets with 20 ground truth clusters.

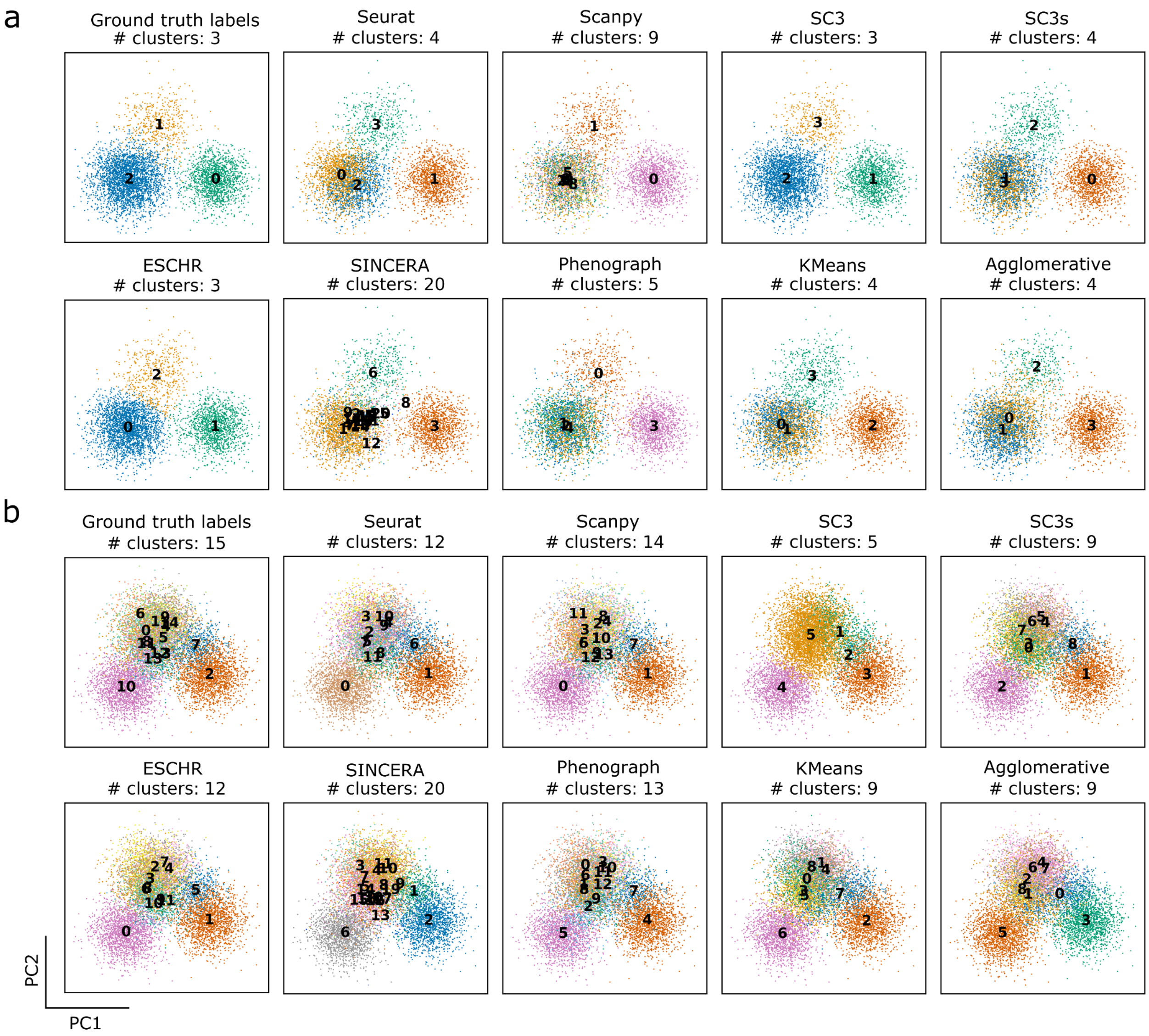

**Supplementary Figure 5: PCA embeddings for example synthetic datasets.** Visualization of data points projected onto the first two principal components, with points colored by either ground truth labels (top left for each panel) or hard cluster labels from the method indicated in the respective title for the remaining 9 plots. (A) shows the dataset used in Figure 2 A-D and (B) shows the dataset used in Figure 2 E-H.

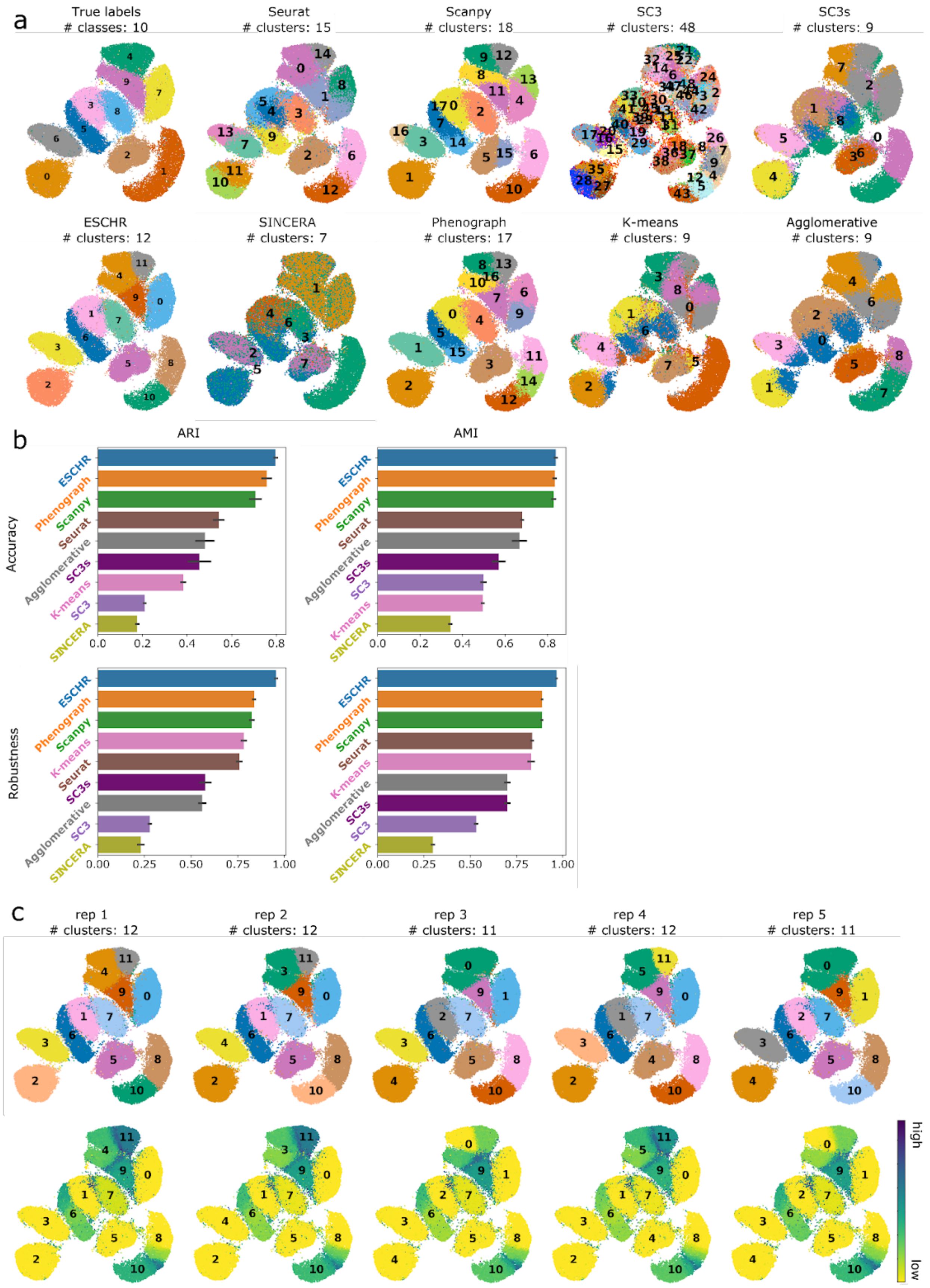
a
True labels
### classes: 10
Seurat
### clusters: 15
Scanpy
### clusters: 18
SC3
### clusters: 48
SC3s
### clusters: 9
ESCHR
### clusters: 12
SINCERA
### clusters: 7
Phenograph
### clusters: 17
K-means
### clusters: 9
Agglomerative
### clusters: 9
b
ARI
AMI
Accuracy
Robustness
ESCHR
Phenograph
Scanpy
Seurat
Agglomerative
SC3s
K-means
SC3
SINCERA
0.0 0.2 0.4 0.6 0.8
0.00 0.25 0.50 0.75 1.00
c
rep 1
### clusters: 12
rep 2
### clusters: 12
rep 3
### clusters: 11
rep 4
### clusters: 12
rep 5
### clusters: 11
high
low

**Supplementary Figure 6: Benchmarking and robustness analysis of ESCHR clustering of the MNIST dataset.** (A) UMAP visualizations of ground truth cluster labels and hard cluster assignments from the first robustness replicate of ESCHR and each of the comparison methods used in benchmarking analysis. Points are colored by cluster ID. (B) Barplots showing accuracy by ARI (top left), accuracy by AMI (top right), robustness by ARI (bottom left), and robustness by AMI (bottom right) for 5 runs of each method on a randomly sampled 90% of the MNIST dataset. Error bars represent 1 standard deviation. Bars are plotted in rank order with highest mean score at the top. (C) UMAP visualizations of ESCHR clustering replicates of the MNIST dataset, where each replicate was generated on a randomly subsampled 90% of data points. First row points are colored by hard cluster label, second row points are colored by overlap score.

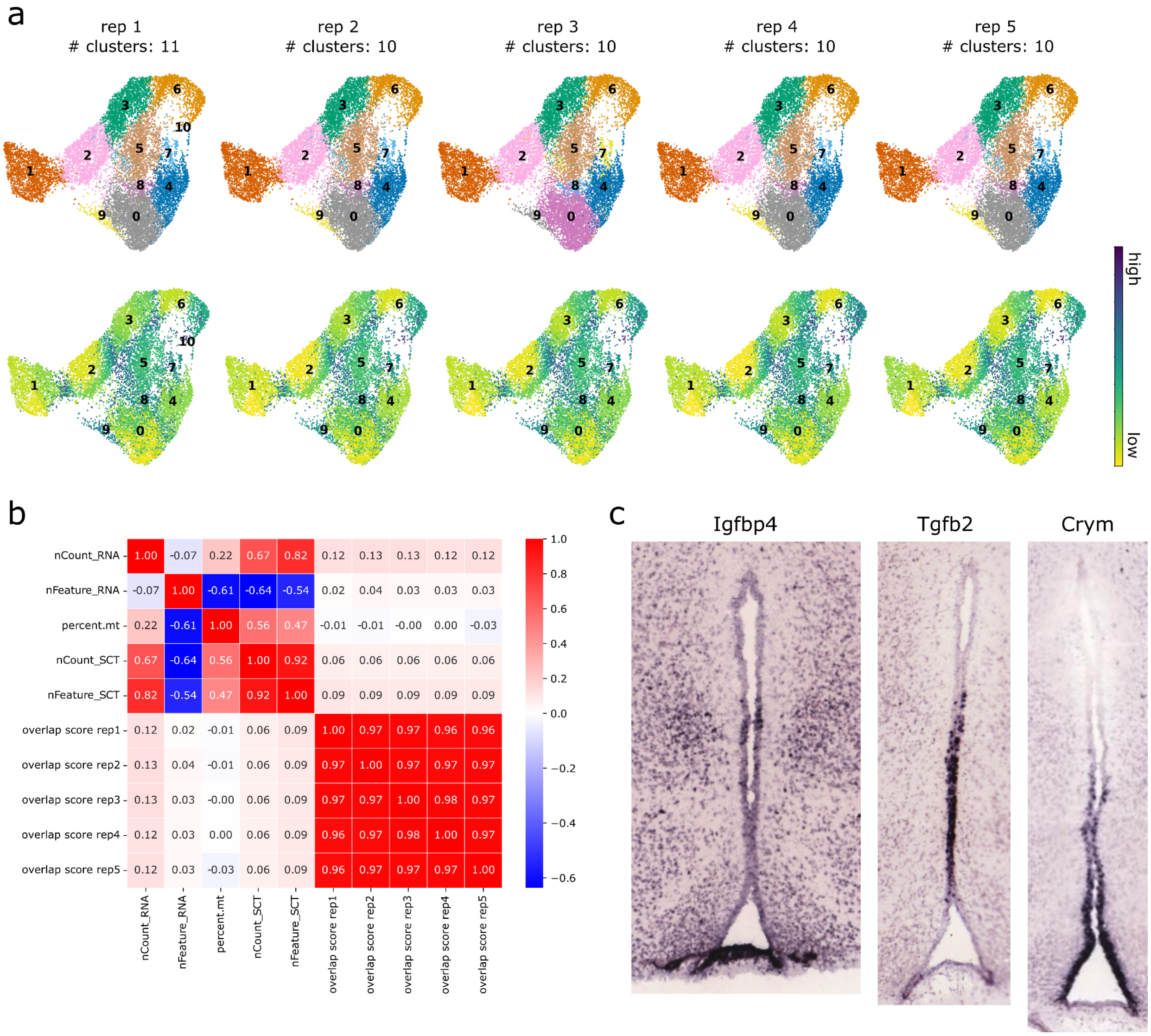

**Supplementary Figure 7: Tanycyte clustering and expression profiles.** (A) UMAP visualizations of ESCHR clustering replicates of Tany-seq data, where each replicate was generated on a randomly subsampled 90% of data points. First row points are colored by hard cluster label, second row points are colored by overlap score. (B) Correlation of metadata features and per-cell overlap scores from each of the 5 robustness replicates visualized in A. (C) Allen Mouse Brain Atlas in situ hybridization images of the full tancyte-containing region.

### Supplementary Tables

**Supplementary Table 1: Overview of methods used in benchmarking analysis.**
*Attached as .xlsx*

**Supplementary Table 2: Overview of real datasets used in benchmarking analysis.**
*Attached as .xlsx*

| method | evaluation | metric | statistic | corrected p-value |
|---|---|---|---|---|
| Seurat | robustness | ARI | 9 | 3.65E-02 |
| Scanpy | robustness | ARI | 1 | 8.50E-03 |
| SC3 | robustness | ARI | 0 | 7.00E-03 |
| SC3s | robustness | ARI | 0 | 7.00E-03 |
| SINCERA | robustness | ARI | 0 | 7.00E-03 |
| Phenograph | robustness | ARI | 0 | 7.00E-03 |
| K-means | robustness | ARI | 0 | 7.00E-03 |
| Agglomerative | robustness | ARI | 0 | 7.00E-03 |
| Seurat | robustness | AMI | 1 | 8.50E-03 |
| Scanpy | robustness | AMI | 0 | 7.00E-03 |
| SC3 | robustness | AMI | 0 | 7.00E-03 |
| SC3s | robustness | AMI | 0 | 7.00E-03 |
| SINCERA | robustness | AMI | 0 | 7.00E-03 |
| Phenograph | robustness | AMI | 0 | 7.00E-03 |
| K-means | robustness | AMI | 0 | 7.00E-03 |
| Agglomerative | robustness | AMI | 0 | 7.00E-03 |
| Seurat | accuracy | ARI | 26 | 4.78E-01 |
| Scanpy | accuracy | ARI | 11 | 5.13E-02 |
| SC3 | accuracy | ARI | 2 | 1.03E-02 |
| SC3s | accuracy | ARI | 0 | 7.00E-03 |
| SINCERA | accuracy | ARI | 0 | 7.00E-03 |
| Phenograph | accuracy | ARI | 0 | 7.00E-03 |
| K-means | accuracy | ARI | 0 | 7.00E-03 |
| Agglomerative | accuracy | ARI | 0 | 7.00E-03 |
| Seurat | accuracy | AMI | 8 | 3.07E-02 |
| Scanpy | accuracy | AMI | 0 | 7.00E-03 |
| SC3 | accuracy | AMI | 3 | 1.24E-02 |
| SC3s | accuracy | AMI | 1 | 8.50E-03 |
| SINCERA | accuracy | AMI | 0 | 7.00E-03 |
| Phenograph | accuracy | AMI | 0 | 7.00E-03 |
| K-means | accuracy | AMI | 0 | 7.00E-03 |
| Agglomerative | accuracy | AMI | 0 | 7.00E-03 |

**Supplementary Table 3: Statistics for benchmarking analysis with synthetic datasets.** Two-sided Wilcoxon signed-rank test with Bonferroni correction was used for statistical analysis comparing ESCHR to each method. N = 20 for comparisons using synthetic datasets.

| method | evaluation | metric | statistic | corrected p-value |
|---|---|---|---|---|
| Seurat | robustness | ARI | 68 | 2.60E-05 |
| Scanpy | robustness | ARI | 49 | 1.38E-07 |
| SC3 | robustness | ARI | 41 | 7.74E-07 |
| SC3s | robustness | ARI | 7 | 8.43E-09 |
| SINCERA | robustness | ARI | 26 | 6.05E-06 |
| Phenograph | robustness | ARI | 97 | 3.31E-05 |
| K-means | robustness | ARI | 38 | 4.89E-08 |
| Agglomerative | robustness | ARI | 37 | 8.55E-06 |
| Seurat | robustness | AMI | 116 | 4.37E-04 |
| Scanpy | robustness | AMI | 105 | 2.71E-06 |
| SC3 | robustness | AMI | 74 | 5.53E-06 |
| SC3s | robustness | AMI | 4 | 7.08E-09 |
| SINCERA | robustness | AMI | 15 | 2.67E-06 |
| Phenograph | robustness | AMI | 197 | 4.76E-03 |
| K-means | robustness | AMI | 41 | 5.77E-08 |
| Agglomerative | robustness | AMI | 39 | 9.83E-06 |
| Seurat | accuracy | ARI | 231 | 1.56E-01 |
| Scanpy | accuracy | ARI | 282 | 1.62E-02 |
| SC3 | accuracy | ARI | 92 | 2.50E-05 |
| SC3s | accuracy | ARI | 572 | 1.00E+00 |
| SINCERA | accuracy | ARI | 127 | 6.62E-03 |
| Phenograph | accuracy | ARI | 284 | 2.21E-01 |
| K-means | accuracy | ARI | 590 | 1.00E+00 |
| Agglomerative | accuracy | ARI | 340 | 1.00E+00 |
| Seurat | accuracy | AMI | 324 | 1.00E+00 |
| Scanpy | accuracy | AMI | 543 | 1.00E+00 |
| SC3 | accuracy | AMI | 110 | 6.80E-05 |
| SC3s | accuracy | AMI | 457 | 1.00E+00 |
| SINCERA | accuracy | AMI | 52 | 6.20E-05 |
| Phenograph | accuracy | AMI | 453 | 1.00E+00 |
| K-means | accuracy | AMI | 524 | 1.00E+00 |
| Agglomerative | accuracy | AMI | 376 | 1.00E+00 |

**Supplementary Table 4: Statistics for benchmarking analysis with real datasets.**
Two-sided Wilcoxon signed-rank test with Bonferroni correction was used for statistical analysis comparing ESCHR to each method. N = 42 for comparisons using real datasets.
